# Germ plasm localisation dynamics mark distinct phases of transcriptional and post-transcriptional regulation control in primordial germ cells

**DOI:** 10.1101/2020.01.12.903336

**Authors:** Fabio M. D’Orazio, Piotr Balwierz, Yixuan Guo, Benjamín Hernández-Rodríguez, Aleksandra Jasiulewicz, Juan M. Vaquerizas, Bradley Cairns, Boris Lenhard, Ferenc Müller

## Abstract

In many animal models, primordial germ cell (PGC) development depends on maternally-deposited germ plasm to avoid somatic cell fate. Here, we show that PGCs respond to regulatory information from the germ plasm in two distinct phases and mechanisms in zebrafish. We show that PGCs commence zygotic genome activation together with the rest of the embryo with no demonstrable differences in transcriptional and chromatin accessibility levels. Thus, cytoplasmic germ plasm determinants only affect post-transcriptional stabilisation of RNAs to diverge transcriptome from somatic cells, which, unexpectedly, also activate germ cell-specific genes. Perinuclear relocalisation of germ plasm is coupled to dramatic divergence in chromatin opening and transcriptome from somatic cells characterised by PGC-specific chromatin topology. Furthermore, we reveal Tdrd7, regulator of germ plasm localisation, as crucial determinant of germ fate acquisition.

## INTRODUCTION

In sexually-reproducing organisms, the germ line ensures that parental genetic information is passed on from one generation to the next. The commitment of the embryonic germ line follows unique steps (Strome and Updike, 2015): in metazoans, the germ fate can be either oocyte-inherited (predetermined) (Eddy, 1975; Williamson and Lehmann, 1996) or zygotically-triggered (induced) (Lawson et al., 1999; Ying et al., 2001). In mammals, germ cells are generated during gastrulation in response to extracellular signals from the surrounding embryonic cells (Ying et al., 2001). On the other hand, most non-mammalian model organisms, such as *C. elegans, D. melanogaster, X. laevis and D. rerio,* require maternal transmission of germ cell-specific factors, which aggregate in a self-containing structure known as germ plasm and distribute into primordial germ cells (Eddy, 1975; Seydoux and Braun, 2006). The germ plasm has been shown to be sufficient and necessary to trigger the germ fate (Gross-Thebing et al., 2017; Tada et al., 2012). The function of the germ plasm is to avoid somatic lineage differentiation of the host cells by at least two mechanisms. Firstly, germ plasm components post-transcriptionally regulate maternal RNA stability and translation (Charlesworth et al., 2006; Iguchi et al., 2006; Nakamura et al., 2004; Siddall et al., 2006; Wilhelm et al., 2003) while they are cleared in the rest of the embryo (Giraldez et al., 2006; Mishima et al., 2006). Also, germ plasm-specific factors, such as Nanos, Dazl and Dead-end (Dnd) function in RNA processing pathways and are indispensable for PGC development (Köprunner et al., 2001; Oulhen et al., 2017; Suzuki et al., 2010). Dnd, induces selective translation of mRNAs by liberating them from micro-RNA (miRNA) inhibition (Kedde et al., 2007), while the RNA-binding-Protein (RBP) DAZL was found to inhibit the translation of several mRNA involved in pluripotency, somatic differentiation and apoptosis in murine PGCs (Chen et al., 2014). Secondly, germ plasm components have been associated with block or delay of Zygotic Genome Activation (ZGA) of the hosting cell and proposed as key factors for avoiding somatic cell fate. In *C. elegans* and *D. melanogaster,* the germ plasm proteins PIE-1 and Pgc delay ZGA in the PGCs, allowing the disengagement between the germ and the somatic lines (Batchelder et al., 1999; Mello et al., 1996; Strome and Updike, 2015).

In contrast to the extensive, genome-wide DNA demethylation observed in migrating mammalian PGCs (Bender et al., 2004; Gkountela et al., 2015; Guo et al., 2015; Tang et al., 2015) reprogramming of DNA methylation has not been seen in zebrafish (Ortega-Recalde et al., 2019; Skvortsova et al., 2019). On the other hand, epigenetic regulators maternally transmitted via germ granules have been implicated in germ fate acquisition (Gaydos et al., 2012; Rechtsteiner et al., 2010; Strome et al., 2014), suggesting that alternative mechanisms of germ plasm-mediated transcriptional regulation may exist.

In this study, we aimed to characterise the potential roles of the germ plasm during PGC formation. We hypothesized that the distinct localisation patterns of the germ plasm before and during PGC migration may represent distinguishable cytoplasmic and nuclear functions in PGC specification. We profiled transcriptome, open chromatin and DNA methylome of developing PGCs at high temporal resolution and discovered two distinct phases of PGC specification during zebrafish embryogenesis. Consequently, we suggest that the first function of the germ plasm is solely cytoplasmic and does not influence transcription or chromatin landscape of the pre-migrating PGCs. However, the second phase of germ cell formation requires chromatin reorganisation, resulting in extensive transcriptional changes that temporally coincide with the association between germ plasm and nuclear membrane. By systematic identification of open chromatin and subsequent prediction of cis-regulatory elements, we show that putative enhancers in proximity of developmental genes remain compacted during PGC specification, resulting in the lack of somatic gene expression. Moreover, predicted PGC-specific regulatory elements appear to be distinctly proximal to the Transcription Start Site (TSS), in sharp contrast to the somatic-specific elements, which spread more distally. The observed chromatin organisation of regulatory elements may underlie unique DNA topology contributing to repression of somatic lineage differentiation. Finally, by inhibiting the translation of Tudor Domain 7a (Tdrd7a), which leads to disruption of germ plasm perinuclear localisation, we demonstrate the requirement of this germ cell determinant in defining germ cell-specific open chromatin and transcriptional landscape.

## RESULTS

### Characterisation of primordial germ cell transcriptome before and during migration

In order to investigate the roles of germ granules, we aimed to characterise the early phases of germ line development via extensive profiling of epigenetic and transcriptional features. For this reason, we focused on the first day of zebrafish embryogenesis, when the PGCs show the most dynamic and active behaviour. The Tg(Buc-GFP) line of *D. rerio* with fluorescently-marked germ plasm (Riemer et al., 2015) was utilised to isolate PGCs and non-fluorescent somatic cells by FACS (Supplementary Figure 1A). Total transcriptome, open chromatin and DNA methylation were analysed at various stages of zebrafish PGC development (Figure 1A). We first assessed transcriptome features associated with developmental stages and cell type and identified major changes coinciding with key events of development. As shown in Figure 1B and C, hierarchical clustering and Principal Component Analyses (PCA) demonstrated minimal transcriptome variance at and immediately after zygotic genome activation (high and dome stages) with small observable difference between replicates of germ plasm-containing and somatic cells (Figure 1C, Supplementary Figure 1B and Supplementary Table 1). However, gradual divergence between the somatic and PGC transcriptomes was coincidental with migration of PGCs and perinuclear localisation of the germ plasm (at 10 somites stage), leading to a marked separation of steady-state transcript content between PGCs and somatic cells by prim-5 stage.

**Figure 1.**
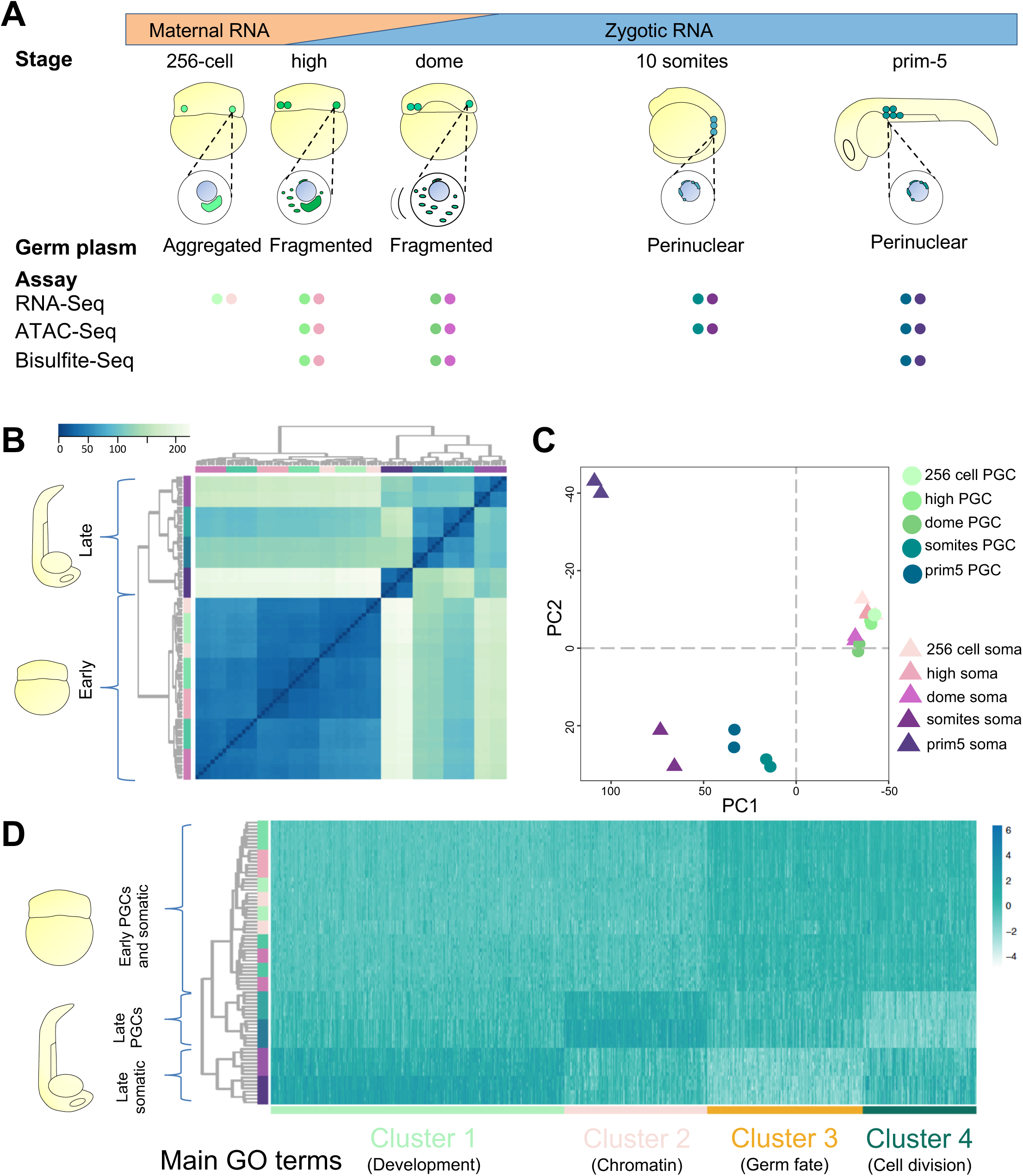
Characterisation of PGC transcriptome highlights early developmental similarities and late divergence between PGCs and somatic cells. (A) Key developmental stages used in the study are shown. Time points were selected according to various phases of germ plasm distribution/PGCs localisation. The early stages span zygotic genome activation including the first wave at 256-cell stage. NGS assays performed for each time point are shown as coloured dots; PGCs and somatic cells are in shades of green and purple respectively. (B-C) Unsupervised hierarchical clustering heatmap and two-dimensional PCA plot show developmental trends of PGC and somatic cell transcriptomes during development. (D) Four clusters of differential gene expression reported as heatmap upon k-mean-based clustering over development and cell type.

In order to determine the transcriptional contribution to PGC development over time, we classified differentially expressed genes into four main groups of temporal expression patterns using k-means clustering (Figure 1D and Supplementary Figure 1C). Within each cluster, genes with distinct biological functions could be identified, as highlighted by group-specific enrichment for gene ontology terms (Supplementary Figure 1D). Genes belonging to Cluster 1 were upregulated in the somatic cells at 10 somites and prim-5 (late) stages over every other sample. As expected, these were associated with developmental processes, differentiation and protein translation. Cluster 2 included those genes upregulated in late PGCs. Interestingly, we noted several pathways involved in chromatin reorganisation, in particular related to DNA packaging. At this developmental stage, germ cells are found in the future genital ridge and show a very low proliferative activity. Accordingly, genes downregulated in the PGCs at late stages versus all other stages were mainly involved in cellular division (Cluster 4), while the Cluster 3 grouped genes for germ fate and thus validate the successful sorting of PGCs.

Taken together, these results demonstrate successful isolation and genome-wide comparative characterisation of PGCs at various developmental stages, which show stage- and cell type-specific transcriptomic differences. Overall, early germ cells have a similar transcriptional profile with the rest of the embryo, which gradually diverges as lineage specification proceeds. At the prim5 stage, analysis of PGC transcriptomes suggests reduced cell proliferation and epigenetic reorganisation compared to somatic cells (Cluster 2 and 4, Supplementary Figure 1D).

### Zebrafish primordial germ cells do not delay zygotic genome activation

Global transcriptomic analysis suggested that PGCs and somatic cells are broadly similar during early zebrafish development. Therefore, although other germ plasm-dependent organisms delay transcriptional activation in the PGCs, the mild transcriptional difference between PGCs and somatic cells at ZGA suggested that transcriptional activation of the germ line may occur alongside the rest of the embryo. We therefore asked whether gene expression in PGCs is repressed during the time when ZGA commences in all the blastomeres. We utilised a recently developed in-vivo transcription imaging tool (MOVIE) and 4D imaged the accumulation and localisation of *microRNA-430* (*miR-430*) primary transcripts, which are the earliest known expressed genes highly active during ZGA in zebrafish (Giraldez et al., 2006; Hadzhiev et al., 2019). Upon injection of fluorescently-labelled morpholinos, we monitored *miR-430* expression in embryos in which germ plasm was tracked by GFP. Interestingly, *miR-430* expression was detectable in somatic cells as well as germ plasm-containing cells already before the main wave of genome activation (Figure 2A, Supplementary Figure 2A and 2B), suggesting that germ plasm does not substantially delay/inhibit early transcription. Also, *miR-430* expression faded around the onset of epiboly in both PGCs and somatic cells, suggesting similar temporal transcriptional regulation between the PGCs and somatic cells.

**Figure 2.**
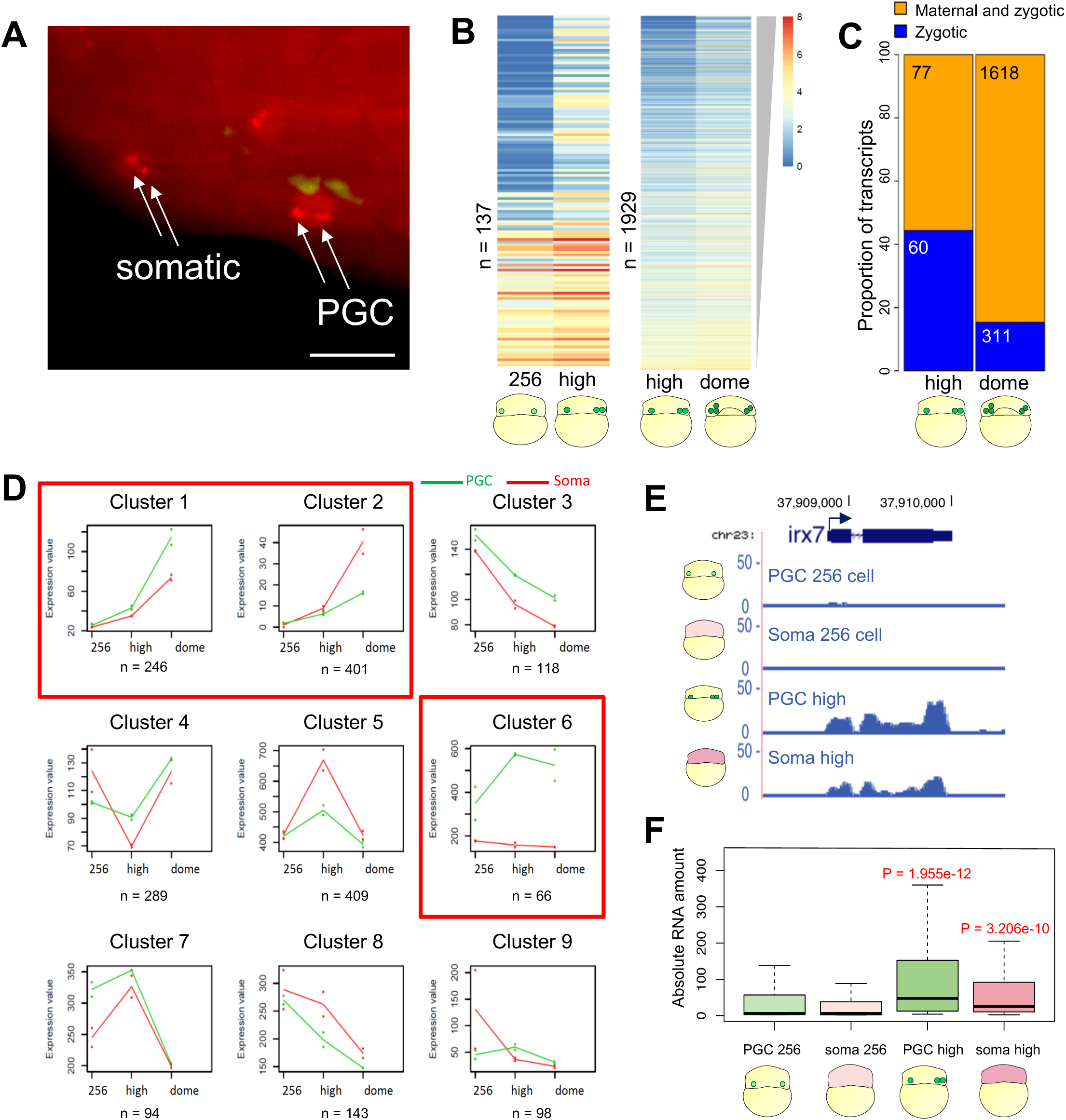
PGCs do not delay the major wave of transcriptional activation. (A) Maximum intensity projection of a multi-stack image showing *miR-430* transcription foci (arrows) detected by fluorescently-tagged Morpholinos (red) in germ plasm-carrying cells and somatic cells at 512-cell stage. Germ plasm-localised GFP-Buc (green) marks PGCs. Scale bar is 30 µm. (B) Heatmap of upregulated genes sorted by log_2_FC in the germ plasm-carrying cells between 256-cell and high stages and high and dome stages. Colours represent the log_2_(tpm+1). P adjusted < 0.1. (C) Proportion of zygotic and maternal/zygotic genes upregulated in the germ plasm-carrying cells at high and dome stages. (D) Clusters of gene expression trends among three developmental stages in PGCs and somatic cells. Median profiles of gene expression values are plotted for each trend in green (PGCs) and red (somatic cells) and represent the gene read counts normalised by the DESeq2-calculated size factor. Red squares highlight clusters supporting transcriptional activation in the PGCs. (E) Genome browser view of normalised RNA-seq reads for *irx7* gene. (F) Absolute RNA concentration normalised to internal control (ERCC spike-in) for genes upregulated in germ plasm-carrying cells between 256-cell and high stage. P-values are calculated by Wilcoxon test.

In order to better understand the relation between ZGA and germ plasm, we studied the composition of the PGC transcriptome by identifying differentially expressed genes across ZGA via RNA-seq (Methods). The number of significantly upregulated genes increased more than an order of magnitude when the germ plasm-carrying cells transition from 256-cell to high stage (135 genes when padj < 0.1 and 124 when padj < 0.05) or from high to dome stages (1926 genes when padj < 0.1 and 1629 when padj < 0.05) (Figure 2B, 2E and Supplementary Table 2). Interestingly, of the 135 upregulated genes after ZGA, 60 were predicted to be zygotically transcribed without maternal contribution (Figure 2C). Zygotic genes were predicted as those with expression lower than 2 tpm at 256-cell stage (no zygotic transcription), which show an increase in expression levels at the subsequent analysed developmental stage. To validate our analysis, we compared our list of predicted zygotic genes with an independent RNA-seq dataset (White et al., 2017). The high degree of overlap between the predicted zygotic and maternal genes within the two datasets supported our arbitrary discrimination between maternally-deposited and zygotically-expressed transcripts (Supplementary Figure 2C). Notably, out of 60 zygotic genes found upregulated in the PGCs at high stage, 56 matched predicted zygotic genes from analysis of the independent RNA-seq experiment (data not shown). This observation suggests that transcription activation does occur at the time when the rest of the blastomeres are also becoming transcriptionally active. Interestingly, among the significantly upregulated genes in the PGCs at high stage, germ cell markers and germ plasm-localised transcripts such as *ddx4*, *dnd1*, *tdrd7a*, *gra* and *dazl* were found upregulated from the previous stage (Supplementary Figure 2D). This was unexpected as germ plasm markers are of maternal origin and are known not to be transcribed until after gastrulation (Blaser et al., 2005; Knaut et al., 2000; Weidinger et al., 1999).

Then, the occurrence of ZGA in the germ plasm-carrying cells was further confirmed after performing a regression-based clustering of genes with similar expression profiles for three stages spanning the first wave of ZGA (256-cell, high and dome) in PGCs and somatic cells. Interestingly, we found two clusters of genes upregulated in both PGCs and somatic cells and one in which genes were upregulated in PGCs exclusively (Figure 2D). Finally, to further verify the observation of occurrence of active transcription between 256-cell and high stage in PGCs, we measured absolute RNA levels over time. After normalisation for an internal RNA control (Jiang et al., 2011) (Supplementary Figure 2F), we found a significant increase in transcript levels in both PGCs and somatic cells at high stage, confirming that active transcription is present in both cell types before the high stage (Figure 2F).

Based on our results, we concluded that the germ plasm does not cause general transcriptional repression in zebrafish, and zebrafish PGCs do not delay ZGA as it has been seen in *C. elegans*, *D. melanogaster* or *X. laevis*.

### Selective retention of zygotically-produced transcripts explains transcriptome differences between germ plasm-carrying cells and somatic cells at ZGA

The observations that germ plasm-carrying cells do not delay transcriptional activation in comparison to the rest of the embryo -yet they appear to carry de-novo generated germ cell-specific transcripts-prompted us to ask whether differential transcription occurs between germ plasm-carrying cells and somatic cells at ZGA. Therefore, we sought to study the nature of ZGA in the pre-migratory PGCs in comparison to the somatic cells. To verify whether the PGCs undergo selective transcriptional activation, we performed differential gene expression analysis of isolated germ plasm-carrying cells and somatic cells at each stage spanning ZGA period. This analysis revealed that before ZGA, already a total of 22 genes were differentially expressed between the two cell types (padj < 0.1) (Figure 3A), confirming that maternal mRNAs are selectively retained in the PGCs as shown previously (Eno et al., 2018; Gazdag et al., 2009; Gerovska and Araúzo-Bravo, 2016; Gorokhova et al., 2007; Levine et al., 2000; Rothschild et al., 2013). However, it was noteworthy that 11 out of 22 identified transcripts have not yet been associated with germ cell or PGC functions and are candidates for novel maternal germ plasm-transcripts. The remaining 11 transcripts instead were either known germ plasm markers (*gra, tdrd7, rg514a, ca15b, dnd1* and *dazl*) or being previously associated with germ cell development/survival (*hook2, tgfa, zswim5, b4galt6, camk2g1*) (Supplementary Table 3).

**Figure 3.**
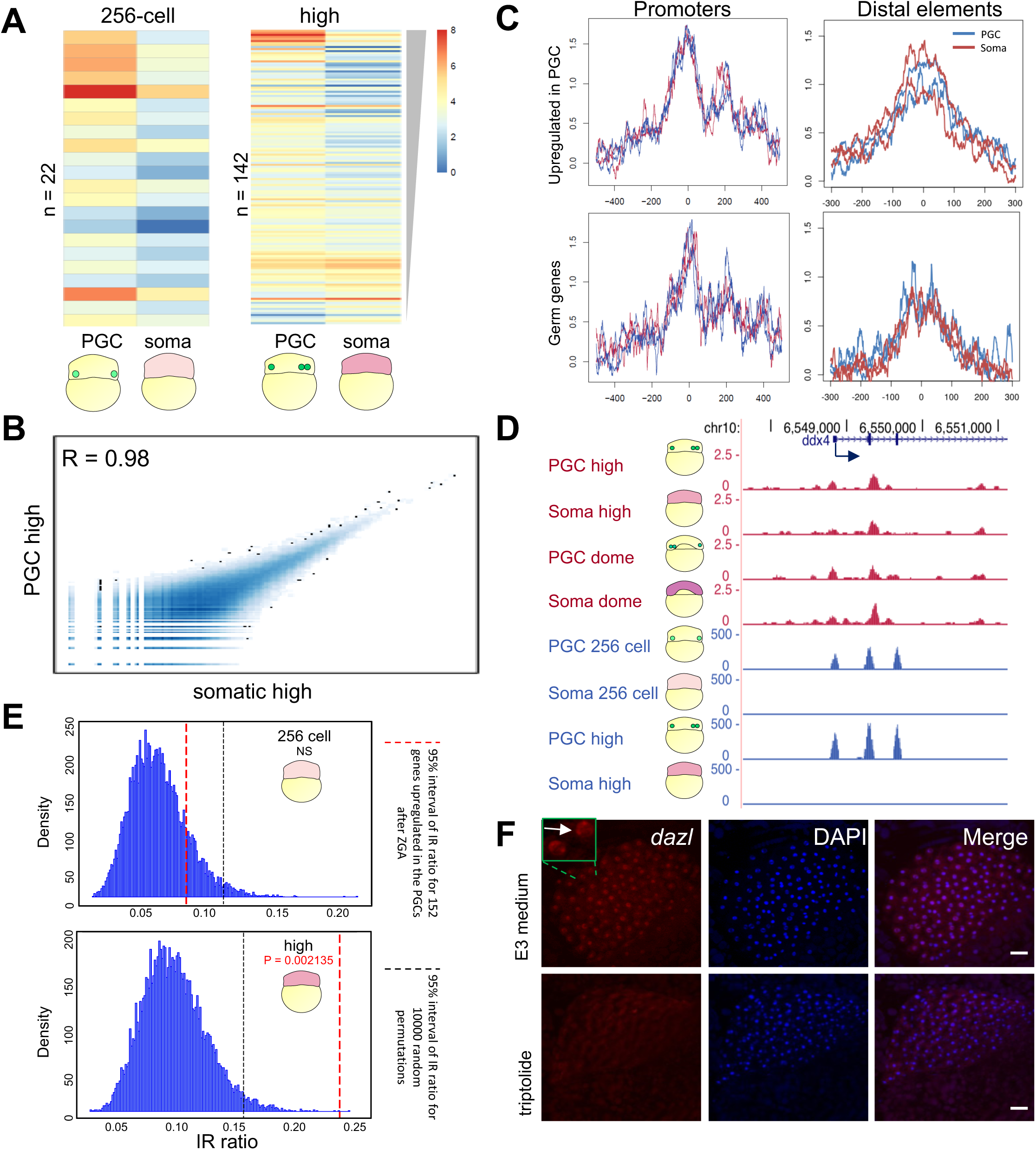
Differential transcriptome between PGCs and somatic cells at early stages is not caused by differential transcription. (A) Heatmap of differentially expressed genes sorted by log_2_FC between PGCs and somatic cells at 256-cell and high stage. Colours represent log_2_(tpm+1) values. P adjusted < 0.05. (B) Correlation plot for open chromatin regions detected in PGCs and somatic cells at high stage. (C) Average accessibility signal at promoters and distal elements of PGC and somatic cells for subgroups of genes at high stage. Promoters are aligned to transcription start site, while distal elements are aligned to peak centre. (D) Genome browser view of normalised ATAC-seq (magenta) and RNA-seq (blue) reads for PGC and somatic cells at stages spanning ZGA. (E) Intron retention ratio before and after ZGA in the somatic cells of genes upregulated in the PGCs after ZGA. P-value is calculated by t-test. Statistical significance was calculated upon 10000 random permutation whose density is shown before and after ZGA. Black and red dashed lines show 95% significance interval for the 10000 permutations and the gene subgroup respectively. NS = not significant. (F) In-situ hybridization for *dazl* pre-mRNA at high stage. Scale bar is 50 µm.

During genome activation (high stage), we identified 142 genes significantly differently regulated between PGCs and somatic cells (Figure 3A). Interestingly, when the number of upregulated genes from one stage to the next was taken into account, we noted a similar trend of gene upregulation in both PGCs and somatic cells, indicating that both the cell types experienced equal transcriptional activation signals (Supplementary Figure 3A). By confirming that ZGA in PGCs and somatic cells is similar, we observed that the relative abundance of transcripts upregulated from 256-cell to high stage in only one cell type was still comparable between germ cells and somatic cells at high stage (Supplementary Figure 3B). Also, the fold change of increased gene expression between 256-cell and high stage showed high correlation between germ plasm-carrying cells and somatic cells (Supplementary Figure 3C). Taken together, these observations suggested that both the cell types activate their genomes similarly.

To address whether retention of the germ plasm in PGCs impacts directly on PGC-specific transcriptional activation and to trace any transcriptional source of the small variation in steady state RNA levels during genome activation, we monitored transcriptional activity using chromatin accessibility state at cis-regulatory elements as a surrogate. We performed Assay for Transposase-Accessible Chromatin combined with sequencing (ATAC-seq) in PGCs and somatic cells. Open chromatin landscapes were analysed globally and compared between PGCs and somatic cells at high and dome stages (start of main ZGA wave). Interestingly, global comparison of open chromatin regions between PGCs and somatic cells at high stage indicated a high degree of correlation (Figure 3B). No distinguishable difference in chromatin accessibility for PGCs and somatic cells was observed on either promoters or distal elements of genes with differential expression between PGCs and somatic cells (Figure 3C). The germ line gene *ddx4*, whose expression was shown by the RNA-seq analysis to increase from 256-cell to high stage only in PGCs, appeared to possess similar degree of chromatin accessibility at its promoter and cis regulatory regions in both cell types (Figure 3D). Taken together, these observations prompted us to hypothesise that, while zygotic transcriptional activation is broadly similar between somatic and germ plasm-carrying cells, the detected differential gene expression was caused by post-transcriptional regulation.

While it was previously reported that maternal RNAs are selectively protected in the PGCs from miRNA-dependent degradation (Mishima et al., 2006), this mechanism has not yet been shown to occur on zygotically-active germ cell genes. As the top-scoring upregulated genes in the PGCs from 256-cell to high stage are known-germ plasm markers, we hypothesised that these are not only deposited by the mother in the germ plasm but are also produced zygotically throughout the early embryo including the somatic cells where they are selectively cleared. To test this hypothesis and to discriminate between maternally-provided and zygotic germ plasm transcripts, we took advantage of our RNA-seq data to perform differential Intron Retention (IR) analysis (Middleton et al., 2017). As intron splicing is co-transcriptional (Merkhofer et al., 2014), newly-transcribed RNAs are expected to show increased intron retention. We compared IR scores for the whole transcriptome before and after ZGA, and observed increase of intron retention in both PGCs and somatic cells upon transcriptional activation (Supplementary Figure 3D). Then we focused on assessing IR in de-novo transcription of germ cell-specific transcripts in somatic cells. A significant increase in intron retention from the previous stage was observed when compared to random sampling of the dataset (Figure 3E). This result further supports the hypothesis that genes transcribed in the PGCs at ZGA were also activated in the somatic cells.

To further validate this observation, we performed quantitative PCR (qPCR) on nuclear and cytoplasmic cell fractions before and after ZGA after removal of PGCs via FACS (Supplementary Table 4). Interestingly, when transcription has started, we saw a significant increase in fold change expression for these genes in the nuclear fraction but not in the cytoplasmic fraction, suggesting that germ cell-related transcripts could be produced by the somatic cells (Supplementary Figure 3E). In order to demonstrate that somatic cells transcribe germ cell-specific RNAs, we studied the localisation of newly-transcribed pre-mRNA within the embryo. Within those transcripts that were selectively upregulated in the PGCs from 256-cell to high stage, *dazl* was one of the highest-scoring hits (Supplementary Figure 3F). We therefore designed RNA probes targeting intronic sequences to visualise unprocessed, newly-produced *dazl* pre-mRNA and carried out fluorescence in-situ hybridization at different stages with and without transcription block. Strikingly, *dazl* was seen to be actively transcribed in the somatic cells, as demonstrated by staining of nuclear foci in the whole embryo at high stage (Figure 3F). Taken together, these results suggest that transcription of germ cell genes occurs throughout the embryo at ZGA. Upon integration with ATAC- and RNA-seq results, we propose that there is no transcriptional delay or differential transcription in germ plasm-carrying cells when zygotic genome is activated. Selective protection of zygotic transcripts together with maternal transcript by the germ plasm may thus contribute to early germ cell specification and onset of migration.

### Primordial germ cells gain specific transcriptomic and epigenetic features during migration

We next asked how transcriptome and chromatin states reflect the distinct ontogeny of PGCs and somatic cells during migration and further development. About 4 hours after fertilisation (dome stage), PGCs initiate independent and guided movements, which will bring them into the future genital ridge site (Raz, 2003). The germ plasm undergoes extensive morphological changes during germ cell migration (Figure 1A). Also, this period coincides with gastrulation and germ layer formation, with remarkable transformation of the whole embryo transcriptome upon lineage diversification (Raj et al., 2018). Our differential gene expression analysis between PGCs and somatic cells indicates increasing divergence of transcriptomes coinciding with germ fate acquisition in the migrating PGCs (as shown in Figure 1D). The number of differentially expressed genes between PGCs and somatic cells gradually increased over time, reaching almost one-third of the total transcriptome at prim-5 stage (Figure 4A). To trace the cell type as the source of transcriptome variation, we looked at the number of upregulated genes from one stage to the in PGCs and somatic cells selectively. We noted that the increase in differential gene expression between PGCs and somatic cells was accompanied by an increase in differential gene expression over time (Figure 4B, Supplementary Figure 4A and Supplementary Table 5), suggesting that both PGCs and somatic cells were undergoing cellular commitment. In-depth analysis of the migratory PGC transcriptome showed overexpression of several chromatin remodelers. The DNA methyltransferase 3 (*dnmt3bb.1*), responsible for de-novo methylation of CpG islands, was upregulated in PGCs when compared to somatic cells, while the Ten Eleven Translocase 2 (*tet2*) was downregulated. Concomitantly, upregulation of the arginine methyltransferase *prmt6*, lysine demethylases (*kdm7* and *kdm8*) and bromodomain-containing proteins (*brdt* and *brd9*) in the PGCs suggested for potential mechanisms of chromatin remodelling via histone modifications (data not shown).

**Figure 4.**
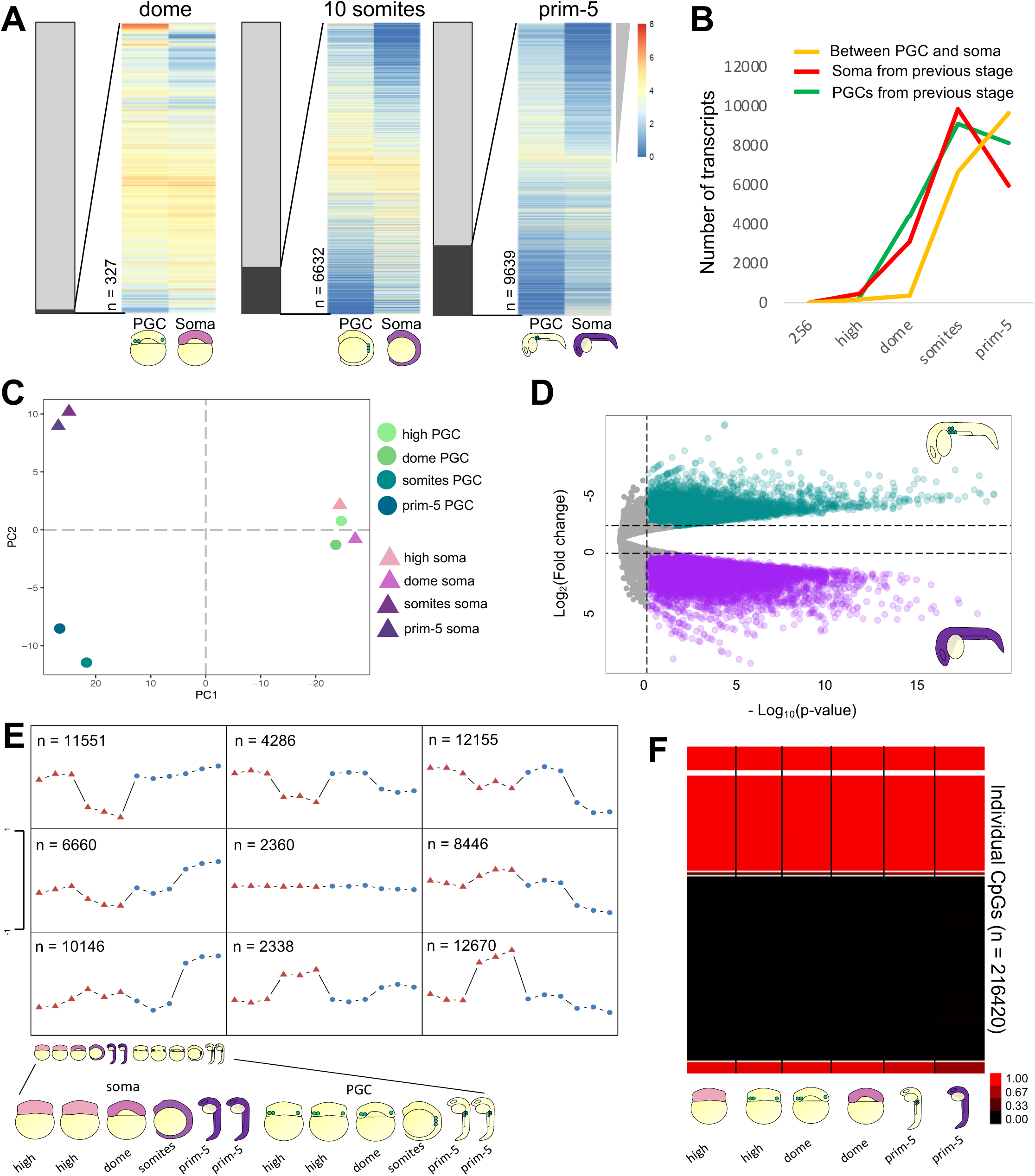
Gradual acquisition of germ identity is accompanied by epigenetic changes. (A) Heatmap of differentially expressed genes sorted by log_2_FC between PGCs and somatic cells. Grey bars represent the proportion of differentially expressed genes over the total transcriptome. Colours represent log_2_(tpm+1) values. (B) Line-chart for gene counts. Upregulated genes from previous stage are in red (somatic) or green (PGCs). The orange line shows number of genes differentially expressed between PGCs and somatic cells at each stage. (C) Two-dimensional PCA plot of ATAC-seq profiles. (D) Volcano plot for regions of open chromatin between PGCs and somatic cells at prim-5 stage (log_2_FC threshold = +/-1, p adjusted < 0.05). (E) Self-organising map of open chromatin regions. PGCs and somatic cells are shown as blue circles and red triangles respectively. Schematic of embryos as in Figure 1A and 1B (F) Methylation status in identified CpGs in PGCs and somatic cells at dome and prim-5 stages.

The differential expression of chromatin regulation prompted us to ask whether germ fate acquisition involves observable epigenetic and chromatin changes as was described in murine PGCs. Therefore, we compared global chromatin accessibility in PGCs and somatic cells via ATAC-seq. After selection of reproducible peaks via irreproducible discovery rate (IDR) filtering (Zhang et al., 2017) (IDR < 0.05) (Supplementary Figure 4B and Supplementary Table 6), we compared global variability of chromatin accessibility among developmental stages and cell types (Figure 4C and Supplementary Figure 4C). In accordance with transcriptome results, early PGCs and somatic cells have similar open chromatin profiles and cluster by stage rather than by cell type. In contrast, marked separation of PGC and somatic cell chromatin accessibility profiles was observed after gastrulation and resulted in the identification of lineage specific sites of open chromatin (Supplementary Figure 4D).

Following differential chromatin accessibility analysis, we found 12591 peaks upregulated in the PGCs (logFC < −1, padj < 0.05) and 23771 peaks upregulated in the somatic cells (logFC > 1, padj < 0.05) at prim-5 stage (Figure 4D). We then used self-organising map (SOM) analysis on PGC and somatic ATAC-seq data at different stages to identify patterns of cis-regulatory element accessibility via an unsupervised approach (Figure 4E). We identified 9 clusters of open chromatin dynamics. Of these, SOM identified 9869 sites that were specific for late stage PGCs and 12654 sites that were less accessible in the PGCs compared to the late stage somatic cells. Interestingly, genes in proximity of somatic-specific ATAC peaks were associated with Gene Ontology (GO) terms for embryonic morphogenesis, tissue formation and embryonic development. These results highlighted that germ fate acquisition requires chromatin regulation.

To gain more insight into the epigenetic specification of PGCs under the control of the germ plasm, we profiled the DNA methylome of pre- and migratory PGCs by performing Reduced Representation Bisulfite Sequencing (RRBS) (Murphy et al., 2018). In contrast to mammals (Guo et al., 2015; Hill et al., 2018), we found no extensive DNA methylation reprogramming of PGCs during these stages (Figure 4G and Supplementary Figure 4F). Analysis of differentially methylated CpGs between PGCs and somatic cells identified 3825 significantly differentially methylated regions (Supplementary Table 7) (only 1.77% of all recovered CpGs from all samples), revealing an overall highly similar methylation programme, in accord with recent studies (Ortega-Recalde et al., 2019; Skvortsova et al., 2019). Based on this result, we concluded that DNA methylation dynamics and chromatin remodelling in zebrafish PGCs were uncoupled.

The finding that post-migratory PGCs show a specific chromatin accessibility landscape in contrast to the early germ plasm-carrying cells suggests that the onset of germ fate acquisition occurs during PGC migration and coincides with the subcellular re-localisation of the germ granules. In addition, differences between PGCs and somatic cells were more marked when chromatin accessibility was profiled in comparison with DNA methylation. This prompted us to investigate more closely the contribution of chromatin regulation to gene expression on different genetic elements.

### Primordial germ cell-specific open chromatin profile is enriched for promoter-proximal putative enhancers and depleted for promoter-distal, developmental putative enhancers

After having established that migrating PGCs undergo chromatin rearrangements, we asked how their epigenetic features contribute to transcription and germ fate acquisition. First, we discriminated between core promoters and distal cis-regulatory elements based on ATAC-seq results in migrating PGCs and somatic cells (prim-5 stage). Gene promoters were defined as sequences within 500bp from annotated Transcription Start Sites (TSSs), while the excluded ATAC peaks (accessible chromatin) within 50kb from the closest TSS were defined as distal elements. Comparison between normalised ATAC signals in PGCs and somatic cells showed that the chromatin profiles were more dissimilar on non-promoter associated open chromatin regions (Figure 5A). In order to further dissect the chromatin accessibility state across the genome, we performed differential chromatin accessibility analysis (logFC > 1, padj < 0.05, fold enrichment > 4) and we focussed on the distribution of differentially regulated ATAC peaks over genic elements. While most of the somatic ATAC peaks occurred at promoter-distal regions, PGC-specific ATAC peaks tended to coincide with promoter (Supplementary Figure 5A). In the PGCs, 43% of the upregulated open regions were found within 1kb from the TSS, while 11% was associated with introns. On the other hand, 41% of the upregulated regions in the somatic cells were found within introns, and only 6% was associated with promoters. We aimed to define putative somatic enhancer regions by intersecting the upregulated ATAC peaks with active enhancer histone mark H3K27ac (Bogdanovic et al., 2012). Out of all identified somatic-specific ATAC peaks (logFC > 1 and padj < 0.05), almost 30% were associated with H3K27ac histone marks. In contrast, less than 10% of PGC-specific ATAC peaks (logFC < −1) matched location of somatic H3K27ac peaks (Supplementary Figure 5B). The functional relevance of cell type-specific chromatin accessibility was estimated by GO analysis of genes associated with differential open chromatin regions away from promoters. As expected, somatic cells were enriched for open chromatin regions in proximity of genes for developmental and differentiation pathways, while PGC-specific ATAC peaks were found in proximity of genes for germ fate, cellular transport and stem cell differentiation (Figure 5B and Supplementary Figure 5C). As GO terms for both gene expression and chromatin accessibility pointed at similar pathways involved in PGC specification, we sought to verify whether accessible DNA regions would be predictive of transcriptional activity. To verify this, we compared the fold change of gene expression between PGCs and somatic cells for a subset of transcripts characterised by significant differential open chromatin between the two cell types. Interestingly, we noted a significant correlation between cell type-specific, promoter-associated ATAC peaks and transcriptional output in both PGCs and somatic cells (Figure 5C, left). On the other hand, the accessibility of distal elements was not predictable of transcription in the PGCs, while a higher correlation between transcription and chromatin accessibility was observed in the somatic cells (Figure 5C, right). This result suggests that transcriptional regulation in migratory PGCs is less dependent on distal elements. In support, we observed an inverse trend of correlation between open chromatin and gene expression in relation to distance from the TSS for PGCs and somatic cells (Supplementary Figure 5D). Moreover, the cumulative distribution of PGC/somatic-specific cis-regulatory elements in relation to the closest transcription start sites showed that PGC-specific cis-regulatory elements were found more proximal to TSSs in comparison to somatic-specific ones (Figure 5D). Taken together, these results indicate that regions of open chromatin predicted to drive gene expression from a distance are less frequent in PCGs.

**Figure 5.**
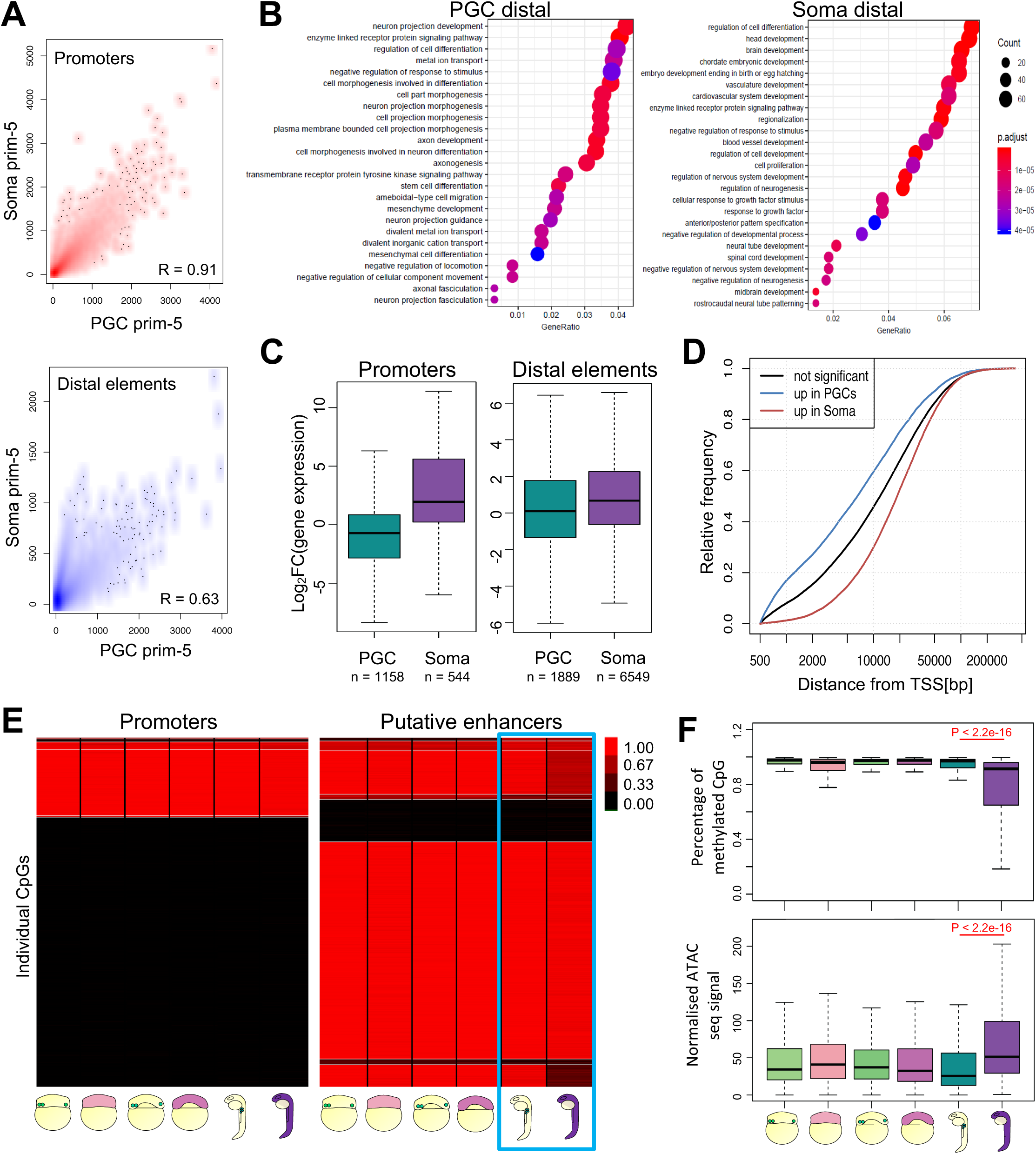
PGCs do not open chromatin at regions identified as putative enhancers. (A) Correlation scatter plots of ATAC peaks between PGCs and somatic cells for promoters and distal elements at prim-5 stage. (B) Developmental processes GO analysis for genes associated with differential chromatin accessible putative enhancers in PGCs and somatic cells. (C) Fold change of gene expression for genes associated with ATAC peaks upregulated in PGCs or somatic cells. Negative and positive fold changes represent gene upregulation in PGCs and somatic cells respectively. (D) Cumulative frequency of open chromatin elements in relation to distance from the closest TSS. P adjusted < 0.05. Colours indicate elements near differentially expressed genes as indicated. (E) Heatmap of DNA methylation for CpGs overlapping promoters (left) and H3K4me1/H3K27ac-rich genomic sites (putative enhancers) in PGCs and somatic cells at high, dome and prim-5 stages. Light blue box highlights putative enhancers at prim-5 stages. (F) Quantification of methylated CpGs and chromatin accessibility at putative enhancer regions. P-value was calculated by Wilcoxon test. Outlier are omitted.

Next we asked what epigenetic mechanisms may be involved in keeping putative distal enhancers closed in PGCs. Two mechanisms have been implicated in enhancer activation: DNA demethylation and histone priming (Bogdanović et al., 2016; Zhang et al., 2018). Although little variation was seen in global DNA methylation profiles of PGCs and somatic cells at prim-5 stage (Figure 4E), we explored methylation dynamics in the PGCs further by discriminating promoter and enhancer regions. When comparing methylation levels across stages and cell types, we observed that promoters preserve their methylation status. Out of 13 validated genes between PGCs and somatic cells, only *ddx4* showed differential DNA methylation on its promoter (Supplementary Figure 5E). In contrast, more dynamic methylation pattern was seen on putative enhancers. In particular, demethylation observed in late somatic cells was not observed in the PGCs, which instead resembled the early profiles characterised by higher levels of DNA methylation (Figure 5E and Supplementary Table 7). Interestingly, putative enhancers showed concomitant higher methylation and significant lower chromatin accessibility in the PGCs compared to the somatic cells (Figure 5F). Next, we tested whether the observed methylation changes were coupled with changes in modulators of DNA methylation. We found upregulation in the PGCs of four out of six *dnmt3bb.1* and *dnmt3bb.2* genes, while no significant change was detected for *dnmt1* gene. On the other hand, *tet2* expression was higher in the somatic cells compared to the PGCs, suggesting that PGCs may lack mechanisms of hydroxymethylation-mediated DNA demethylation catalysis (Breiling and Lyko, 2015; Hill et al., 2018) (Figure 5F).

Based on these results, we propose that PGCs avoid somatic cellular commitment and somatic fate by inhibiting transcription of developmental genes upon block of chromatin opening of their enhancers. Moreover, PGCs appear to transcribe genes which, unlike developmental genes, are enriched in promoter-proximal regulatory elements and thus appear to be regulated by contrasting chromatin topology from that seen in somatic cells.

### TDRD7a, a germ plasm-segregating protein, is required for maintaining germ line-specific chromatin and transcriptome signature

Our genome-wide analysis of chromatin accessibility demonstrated a germ line-specific chromatin reprogramming, which coincides with the transition of the subcellular localisation of the germ granules from cytoplasmic to perinuclear during the early phases of migration (5-6hpf) (Doitsidou et al., 2002; Strasser et al., 2008; Updike et al., 2011; Weidinger et al., 2003). After this stage, the germ plasm associates with the nuclear membrane, although the role of this interaction remains unknown. We hypothesised that the observed transcriptome divergence between PGCs and somatic cells could be driven by perinuclear germ plasm-mediated gene regulation mechanisms detectable on the level of chromatin changes. A known germ plasm marker is the protein Tudor domain 7 (Tdrd7), which is also one of the highest differentially expressed genes in the reported RNA-seq dataset at all developmental stages (data not shown). When translation of Tdrd7 is inhibited by morpholino interference, the germ plasm is incapable of fragmenting and forming small localised granules (Strasser et al., 2008). In order to test whether Tdrd7 and perinuclear germ granules are required for genome-wide chromatin rearrangement observed in PGCs, we investigated the chromatin accessibility and transcriptome profiles of embryos, in which germ plasm localisation was perturbed upon morpholino-mediated Tdrd7 knock-down (KD) (Supplementary Figure 6A and Supplementary Table 8).

Differential gene expression analysis of Tdrd7 KD and wild type PGCs at prim-5 stage indicated a dramatic effect by the loss of Tdrd7 (Figure 6A and Supplementary Table 9). Gene ontology analysis indicated that downregulated genes were enriched for genes involved in reproduction and germ cell development, while upregulated genes were enriched in genes associated with developmental process, organogenesis and protein expression processes (Supplementary Figure 6B). These observations suggested that, in absence of Tdrd7 and correct germ plasm distribution, germ cell character was lost in favour of somatic lineage fate acquisition. Indeed, PCA analysis of variance among Tdrd7 wild type and KD embryo transcriptomes demonstrated that lack of Tdrd7 in PGCs shifted their transcriptome profile towards somatic-like transcriptome (Figure 6B). Detailed RNA-seq analysis highlighted that genes downregulated in PGCs upon Tdrd7 KD also include genes associated with pluripotency and zygotic genes for gametogenesis. While ubiquitously expressed housekeeping genes were unaffected (Figure 6C and Figure 6E). It was noteworthy, that maternally-inherited germ plasm transcripts showed only slightly reduced levels in the Tdrd7-lacking PGCs as compared to control PGCs (Supplementary Figure 6C and Supplementary Figure 6D). This could be caused inhibition of late PGC-specific transcription upon Tdrd7 KD, while maternal mRNAs were preserved. These observations indicate that disruption of Tdrd7 function with germ plasm mis-localisation leads to global deregulation of PGC transcription programme and mild notable effect on germ plasm RNAs.

**Figure 6.**
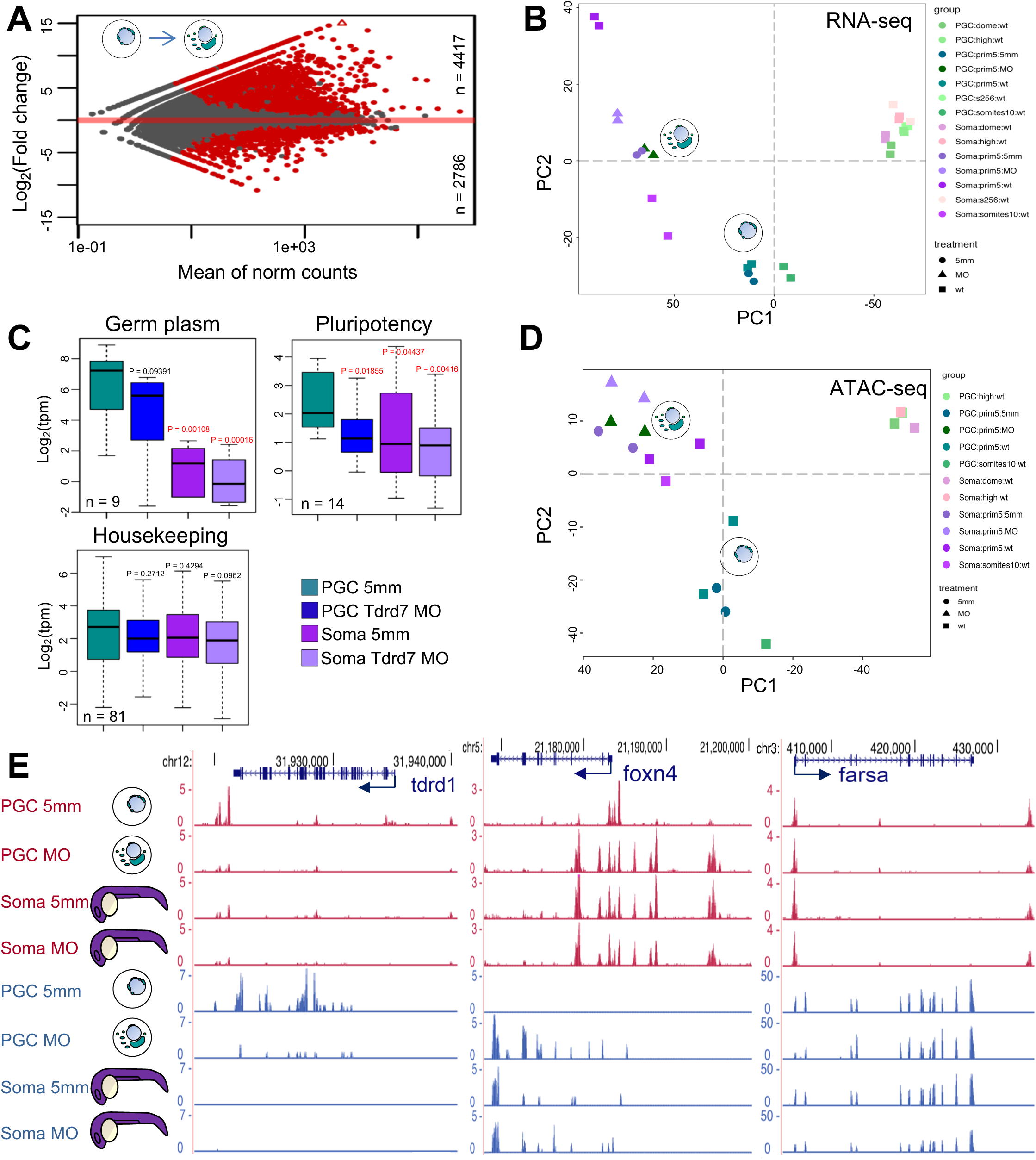
Tdrd7 is required for maintaining PGC fate. (A) Differential expression analysis for genes upregulated and downregulated by Tdrd7 KD in PGCs at prim-5 stage. Significantly differentially expressed genes are in red. P adjusted < 0.1, log_2_FC < −1/> 1. (B) Global transcriptional variance shown as PCA plot for wild type and MO-injected PGCs and somatic cells. (C) Boxplots reporting normalised transcript levels (tpm values) for various gene subsets in MO-injected PGCs and somatic cells. P-values against control is shown. Red colour indicates significance (p < 0.05) based on Wilcoxon test. (D) PCA plots for only promoters and putative enhancers show PGCs diverging towards somatic-like chromatin state. (E) Genome browser view of ATAC profiles after morpholino injections. Open chromatin (ATAC-seq) is shown in magenta and transcript levels (RNA-seq) are shown in blue. Arrows show transcriptional directionality.

Next, we asked whether the causes of differential gene expression in Tdrd7 KD PGCs could be traceable to the chromatin state. To this end, ATAC-seq was carried out on Tdrd7 KD PGCs at prim-5 stage and compared to control PGCs. Global analysis of open chromatin in Tdrd7 KD embryos revealed a marked effect by Tdrd7 loss in the PGCs. Open chromatin of Tdrd7 KD PGCs resembled more to that in the wild type somatic cells than to wild type and PGCs injected with a morpholino control, suggesting that the observed transcriptome phenotype was accompanied by chromatin changes (Figure 6D).

Analysis of putative cis-regulatory element in proximity of misregulated genes in Tdrd7 morphant embryos showed a general tendency towards compaction of PGC-specific genes regions, while development- and morphogenesis-associated genes gained open chromatin peaks in comparison to control PGC cells (Figure 6E and Supplementary Figure 6D). To get a more comprehensive overview of mechanisms governing chromatin regulation in the PGCs and to confirm the association between chromatin accessibility profile and apparent reprogramming of PGCs towards the somatic fates, we have carried out a global analysis of open chromatin behaviour upon Tdrd7 knock down and generated SOM classes of putative regulatory elements (Supplementary Figure 6E). Unsupervised sample clustering confirmed epigenetic differences between wild type and Tdrd7 KD PGCs, indicating distinct shift of PGC chromatin states towards somatic fate, when translation of *tdrd7* was inhibited and the germ granules were mis-localised. Of note, PGC-specific genes, such as *dazl*, which are expressed zygotically as well as inherited maternally in the germ plasm, were shown to be associated with closure of promoter and candidate enhancers despite only mild reduction in their RNA levels (Supplementary Figure 6D).

These results demonstrate transcription regulatory roles for the germ plasm determinant Tdrd7 and suggest that perinuclear germ plasm-nucleus cross-talk is required for germ cell-specific gene activation and fate decision.

## DISCUSSION

In this study, we have investigated the transcriptome, chromatin accessibility and DNA methylation dynamics during early development of PGCs in a germ plasm-dependent vertebrate. The use of the Tg(Buc-GFP) line, combined with ATAC- and RNA-seq, has allowed us to investigate the nuclear and cytoplasmic roles of the germ plasm at unprecedented resolution. Moreover, by following chromatin and transcriptional features in the germ line over time, we were able to describe two distinct roles for germ cell-specific cytoplasmic granules. We link these two roles with two distinct germ plasm subcellular distributions and distinguished an early and a late phase of germ fate acquisition in zebrafish (Figure 7). In contrast to many animal models, we demonstrated that germ plasm contributes to post-transcriptional regulation of both maternal and zygotic gene products, with no detectable effect on transcription during and after genome activation. Following commencement of PGC migration and re-localisation of the germ plasm, PGC-specific chromatin accessibility profiles evolve together with PGC-specific transcription, which we show are Tdrd7-dependent and likely mediated by re-localisation of the germ granules.

**Figure 7.**
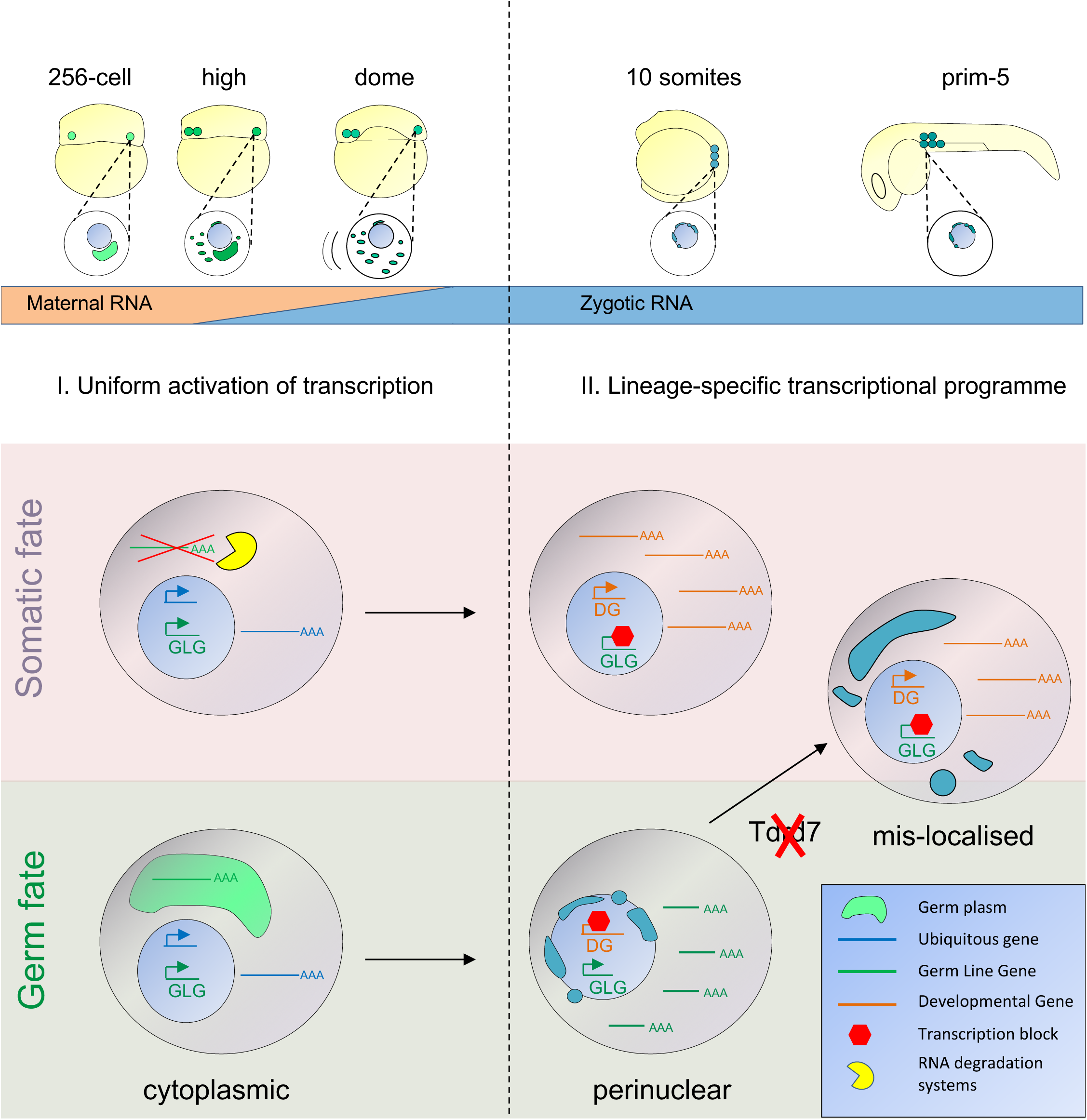
Two functions of the germ plasm during germ fate acquisition. During zygotic genome activation, both somatic cells and PGCs activate the genome similarly, including the activation of PGC specific genes. However, PGC specific genes are only stabilised in germ plasm-containing cells upon protection from the zygotic degradation machinery (yellow). During PGC migration, germ plasm components re-localise to the nuclear membrane. This coincides with acquisition of a PGC-specific transcriptional programme, involving transcriptional repression of developmental pathways blocking somatic differentiation. This process is Tdrd7 dependent as indicated by loss of germ plasm re-localisation and reactivation of somatic transcriptional programmes.

### ZGA is not delayed in zebrafish germ plasm-carrying cells

Germ plasm factors block or delay zygotic genome activation by sequestering RNA polymerase II (RNA Poll II) or its co-factors at the time of transcriptional activation in many organisms (Batchelder et al., 1999; Mello et al., 1996). By delaying the ZGA, the PGCs avoid the commencement of developmental differentiation programmes characteristic to the developing somatic lineages. In mouse, in which PGCs develop by induction and without maternal germ plasm (Lawson et al., 1999; Ohinata et al., 2009; Saitou and Yamaji, 2012), newly-formed germ cells undergo transcriptional quiescence for a short period of time (Kurimoto et al., 2008). The mechanisms of transcription pausing are not fully understood, however they involve inhibition of the elongation factor P-TEFb by Pgc and PIE-1 proteins in *D. melanogaster and C. elegans* (Hanyu-Nakamura et al., 2008; Mello et al., 1996) and sequestration of the general transcription factor TAF4 in *C. elegans* (Guven-Ozkan et al., 2008). In sharp contrast, immunostaining of Serine 2-phosphorylated RNA Poll II, showed localised nuclear foci in zebrafish germ plasm-carrying cells as early as 256-cell stage (Knaut et al., 2000), suggesting that delay of genome activation in the germ line may not occur in this species and that alternative mechanisms of germ fate acquisition may exist. In this study, we imaged transcription in-vivo for the first time in PGCs and together with transcriptome analysis, provide independent lines of evidence that the germ plasm does not delay the first wave of Zygotic Genome Activation (ZGA) in zebrafish. These findings indicate striking plasticity in establishing the germ cell fate among clades and raise the prospect that transcriptional delay is not essential for PGC formation. Interestingly, since massive transcriptional activation has been previously shown to cause double strand break (DSB) in the embryonic germ line (Butuči et al., 2015), by not delaying ZGA zebrafish PGCs may experience a milder transition from a non-transcribing to a transcribing state while being less exposed to DNA damage.

### Why do PGCs need zygotic contribution to maternally deposited PGC mRNAs?

We provide several lines of evidence to show that PGCs and somatic cells transcribe similar set of genes during blastula stages and that there is no overt difference in their chromatin states. We analysed chromatin accessibility, de-novo transcript abundance, intron retention and intron-containing nascent RNA localisation, which all indicate no difference in transcription between the PGCs and the somatic cells. Strikingly, we report that PGC-specific RNAs are also zygotically transcribed in the somatic precursors, where their maternal counterparts are known to be promptly degraded by zygotic machineries (Giraldez et al., 2006; Mishima et al., 2006). Thus, it is unclear why somatic cells produce PGC-specific RNAs. This unexpected observation may be explained by chromatin accessibility organisation of undifferentiated pluripotent stem cells, such as the blastomeres of the zebrafish embryo at and immediately after ZGA. Pluripotent cells display less compact chromatin organisation than differentiating cells (Andrey and Mundlos, 2017) and are characterised by accessible chromatin with low, but detectable activity of a broad range of genes. In contrast, differentiating cells are characterised by gradual divergence of accessibility of lineage and cell type specific-enhancers (Ladstätter and Tachibana, 2019; Lu et al., 2016; Perino and Veenstra, 2016). Therefore, it is feasible that during ZGA, zebrafish blastomeres carry a pluripotent ES-like chromatin and corresponding transcription state characterised by broad capacity of gene expression. This primordial state of chromatin organisation, likely lacks or only commences the formation of chromatin architecture characteristic of differentiating cells during development (Kaaij et al., 2018). It is conceivable that, in this primordial chromatin architecture, sophisticated, enhancer-dependent regulation has not yet been established and gene expression occurs without lineage-specificity and as such it tolerates, for example, germ-cell specific gene expression in the somatic progenitors. In this model, emphasis is on post-transcriptional control of gene expression and germ plasm-mediated, selective stabilisation of zygotic transcripts could be the primary source of divergence of transcriptome between PGCs and somatic cells (Figure 7). In support of this, several germ plasm factors are known to contribute to post-transcriptional and translational regulation. For example, the RNA binding germ plasm component Dnd1 stabilises maternal mRNAs selectively by restricting access of miRNAs responsible for clearing maternal mRNAs in the somatic cells (Giraldez et al., 2006; Kedde et al., 2007). Likewise, the germ factor Nanos has been implicated in destabilisation of RNAs and translational inhibition in the germ cells in combination with Pumilio and CCR4-NOT (Lee et al., 2017; Suzuki et al., 2012).

An additional question emerging from our finding is why zebrafish PGCs need zygotic contribution to PGC-specific mRNA activity at ZGA, while most of these PGC-specific RNAs are already present either in the germ plasm or in the cytoplasm of germ plasm-containing PGCs. While we cannot answer this question directly, it is likely that such early activation of zygotic transcription is either redundant, similarly to the thousands of mRNAs which are both present maternally and zygotically in the somatic progenitors (Haberle et al., 2014; Harvey et al., 2013) or that the zygotic component is required to gradually take over and compensate for loss and or dilution of maternal mRNA in dividing PGCs.

### A germ plasm-mediated epigenetic reprogramming engages germ fate

Germ plasm-carrying cells initiate independent and active migratory movements by dome stage (Blaser et al., 2005; Bontems et al., 2009; Eno and Pelegri, 2016; Eno et al., 2018; Raz, 2003; Yoon et al., 1997), suggesting that migration may, in part, be transcriptionally regulated. However, given the minimal transcriptional differences observed between PGCs and somatic cells, the main mode of regulation is expected to be post-transcriptional. As shown before, early migratory movements are triggered by Dnd-mediated loss of cell adhesion (Blaser et al., 2005) as well as the interaction between the C-X-C chemokine receptor type 4b (Cxcr4b) on the PGC surface and the ligand Cxcl12a (Doitsidou et al., 2002). Additionally, translational inhibition of *nanos* and *oskar* mediated by the germ plasm factors Bruno and CUP is required during germ cell formation in *D. melanogaster* (Nakamura et al., 2004; Wilhelm et al., 2003), while the RNA-binding protein DAZL inhibits translation of several mRNAs involved in pluripotency, somatic differentiation and apoptosis in mouse PGCs (Chen et al., 2014). It is thus not surprising that the germ plasm-specific factor *dazl* is degraded after ZGA in the somatic precursors to restrict its function onto the PGCs in zebrafish. While we have not explored the mRNA clearance process involved in removing zygotically-transcribed mRNAs from the somatic cells, this is likely achieved via the global regulator of maternal mRNA degradation, *miR-430* (Giraldez et al., 2006). Based on our results, we speculate that the germ plasm is exclusively required to protect and process both maternal and zygotic transcripts during onset of migration in the germ plasm-carrying cells. This feature provides the germ plasm-carrying cells with unique capacities, likely required for triggering migration and providing foundation for downstream molecular cues, which will initiate the second phase of PGC development.

We have demonstrated, that unique chromatin accessibility programming occurs in the zebrafish PGCs and that this cis-regulatory element-associated topology contributes to avoiding somatic differentiation. Also, we were able to associate germ plasm re-localisation as an extranuclear effector, which acts on chromatin remodelling coinciding with perinuclear localisation (Figure 7). In line with this observation, interference with germ plasm localisation via its regulator Tdrd7 impacts on the PGC fate, which correctly migrate, but experience somatic differentiation during and following migration.

Notably, our results with DNA methylation analysis demonstrate that even in the absence of global DNA demethylation (Macleod et al., 1999; Bogdanović et al., 2016; Skvortsova et al., 2019), local, differential methylation between PGCs and somatic cells is observed at regulatory sites. However, in contrast to mammalian PGCs, where DNA methylation is almost completely erased genome-wide (Guo et al., 2015; Hill et al., 2018), we observe an inverse trend in zebrafish, where hypermethylation is found on somatic putative enhancers, suggesting remarkably different mechanism for epigenetic programming of the germ line among vertebrates. This was further corroborated with open chromatin analysis genome-wide, which showed lack of opening somatic enhancers in PGCs. Interestingly, although our study led to the identification of PGC-specific putative regulatory elements, we found low correlation between opening of distal ATAC peaks and transcription. Higher correlation between chromatin accessibility and transcription was detected for TSS-proximal cis-acting elements in PGCs than in somatic cells. This finding, in line with previous reports in humans (Guo et al., 2017), suggests that transcription activation in PGCs is achieved by short-range interactions. Nevertheless, additional studies focusing on the DNA interactions, Topological Associated Domains (TADs) formation and spatial organisation of the chromatin in the developing germ line will contribute to better understand the chromatin architecture-associated mechanisms of acquisition and maintenance of totipotency.

### Importance of germ plasm components and subcellular organisation

In accordance with previous studies, our results confirm that removal of individual germ plasm components is sufficient to trigger somatic differentiation in the PGCs (Gross-Thebing et al., 2017; Lee et al., 2017). Tdrd7-depleted PGCs preserve germ granules, germ factors and correctly reach the genital ridge (Hosokawa et al., 2007; Strasser et al., 2008), suggesting that pathways upstream migratory movements are unaffected. However, despite this mild phenotype, we report a remarkable somatic-like chromatin accessibility and consequent transcriptional reprogramming. Based on this result, we speculate that lack of Tdrd7 and incorrect germ granule localisation are sufficient to diverge the fate of the embryonic germ line and induce activation of somatic differentiation pathways.

Tdrd7 is known to interact with Piwi, a piRNAs processor (Huang et al., 2011). Piwi-mediated piRNAs processing has been associated with epigenetic changes in both the somatic and the germ line (Houwing et al., 2007), therefore it is tempting to speculate about a link between piRNA pathways and germ fate maintenance. For example, piRNAs are known to control transposon silencing via H3K9me3 and, in general, to regulate chromatin state on piRNA-target regions (Huang et al., 2013; Sienski et al., 2012). Interestingly, it has been recently reported that germ granules protect germ line transcripts from piRNA-mediated silencing, regulating the pace of release from the cytoplasm to the nucleus (Ouyang et al., 2019). To date, we cannot conclude on whether the germ granules re-localisation is epistatic to the epigenetic germ fate acquisition or vice-versa. However, the multiple lines of evidence associating Piwi and piRNAs to epigenetic regulation strongly suggest a possible requirement of the germ plasm-nuclear interaction in order to trigger initiation of epigenetic germ fate. In conclusion, we suggest that perinuclear localisation of the germ plasm and Tdrd7 are involved in chromatin reprogramming of gonadal PGCs during somitogenesis. Our discoveries have fundamental implications in the understanding of pluripotent fate acquisition and the functional relationship between subcellular aggregates with epigenetic and chromatin reprogramming.

## ACKNOWLEDGEMENT

This work was supported by the H2020 programme Zencode-ITN, Wellcome Trust Investigator award (106955/Z/15/Z) and BBSRC (BB/L010488/1) to F.M. and B.L. and HFSP programme grant to F.M. and B.C. and NEURAM FET H2020 project of the European Commission to FM. We thank Brian Dalley for sequencing expertise, K.T. Varley for helping us adapt Patch Bisulfite PCR, the HCI high-throughput genomics core, flow cytometry core, HCI fish facility, and the Centralized Zebrafish Animal Resource (CZAR) facility at the University of Utah. Financial support was received from the Howard Hughes Medical Institute and the Huntsman Cancer Institute core facilities (CA042014). We thank Erez Raz and Katsiaryna Tarbashevicz at the Center for Molecular Biology of Inflammation (ZMBE) in Muenster for advice on the isolation of PGCs and comments on the manuscript. We thank Roland Dosch at the University Medical Center in Göttingen for providing the Tg(Buc-GFP) line and the BMSU facility at the University of Birmingham for zebrafish maintenance. We thank the Tech Hub core at the University of Birmingham for the flow cytometry and NGS equipment and technical support provided.

## AUTHORS CONTRIBUTIONS

F.M.D. and F.M. conceived the study and wrote the manuscript. All the authors critically revised the manuscript. All experiments were performed by F.M.D. except for the following: Y.G. generated RRBS libraries from isolated genomic DNA. A.J. produced light sheet images of morpholino-injected Buc-GFP embryos. ATAC-seq data were analysed by F.M.D. and P.B. RNA-seq analysis was performed by F.M.D. and B.H.R. RRB-seq analysis was designed by B.C. and Y.G and performed by Y.G. and B.C.

## METHODS

### Animal procedures

All animal work was performed under the Project Licence # b6b8b391, in accordance with the UK Home Office regulations and UK Animals (Scientific Procedures) Act 1986. Fish pairs were crossed in 1 litre breeding tanks and kept overnight separated. The next morning, the gate was removed and the eggs collected at intervals of 5 minutes to ensure synchrony. Fertilised eggs were dechorionated by 10mg/mL Pronase and serial washes in sterile E3 medium. After dechorionation, embryos were kept in agarose-coated petri dishes at 28.5 °C in a 14/10 hours of light/dark respectively. Wild type (AB), Tg(Buc-GFP) and Tg(kop:eGFP) lines were used during this work.

### Microinjections of zebrafish embryos

Morpholinos were diluted in phenol red and about 0.3 pM were injected into the yolk of fertilised zebrafish embryos with a glass needle as described in Hadzhiev et al., 2019.

### Transcription block

Transcription block was achieved through embryo incubation in 1 µM triptolide (Sigma T3652) in E3 medium from the single-cell stage to completion of the experiment.

### PGC preparation for FACS

PGCs were isolated at different stages via FACS. Tg(Buc-GFP) heterozygous embryos were grown at the desired stage by incubation in E3 medium supplemented with 1mg/ml gentamicin at 28.5°C. Embryos were washed three times in sterile water before collection and about 200 of them were pulled in single microcentrifuge tubes. 500 µl of HBSS supplemented with 0.25% BSA and 10mM Hepes were added and dissociation occurred by pipetting for 2 minutes with a glass pipette. Excess of yolk was removed by two rounds of 3 minutes centrifugation at 350 x g at 4°C, while pelleted cells were resuspended in 1ml HBSS supplemented with 0.25% BSA and 10mM Hepes prior of filtering. Cell suspension was kept on ice for the entire isolation procedure.

### Fluorescent in-situ hybridization

Dechorionated embryos were collected at the desired stage, washed in cold PBS and fixed in 4% PFA at 4°C for 1 hour. Fixed embryos were then dehydrated in increasing dilutions of methanol (25, 50, 75, 100 %) and left from overnight to one month at −20°C. The embryos were rehydrated in decreasing dilutions of methanol (75, 50, 25%) and washed 5 times in PBST (0.1%) for 5 minutes with gentle agitation. In order to acclimatise the sample to the high temperature and the hybridization conditions, the embryos were incubated for 2 hours at 70°C in 200 µl of Hybridization Buffer (HB) (50% deionized formamide, 5X SSC, 0.1% Tween-20, 50 μg/ml of heparin bile salts, 500 μg/ml of extracted RNase-free tRNA, pH 6.0). From 50 to 100 ng of DIG-labelled RNA probes targeting *dazl* transcripts (Forward: ACTAAAGTTGTAGCTGGGCCT, Reverse: CCTGAGTGGGCGTTAATGTT) were added to the HB and incubated overnight at 70°C. The next day, the probes were removed by four washes in increasing dilutions of 2 X SSC (NaCl 0.3M; Sodium citrate 0.03M) at 70°C (25, 50, 75, 100 %). The sample was then washed twice in 0.2% of SSC at 70°C 30 minutes each. The 0.2 X SSC was replaced by four serial dilutions in PBST 0.1% in order to remove any left-over probe. Washes were performed at room temperature with gentle agitation. The embryos were blocked in Blocking Buffer + Maleic Acid (Roche, 11585762001) for at least 3 hours before the anti-DIG antibody was added in a concentration of 1:5000 and incubated overnight at 4°C. The next day the antibody was washed five times in PBST 0.1% with gentle agitation at room temperature (30 minutes each) and fluorescently-tagged by horseradish peroxidase-catalysed signal amplification (Thermo Scientific, T20913)

### Imaging

Embryos were placed in an agarose-coated petri dish and eventually embedded in agarose and imaged with a Zeiss 780 confocal or Z1 light sheet microscopes. Images were taken with the Zeiss ZEN pro 2.0 acquisition software with standard settings. Fixed samples were mounted in glycerol-based VectaShield (Vector laboratories, H-1000, UK) on a slide and covered with a glass slip. When preservation of the body shape was required, imaging dishes with glass bottom were used to avoid disintegration of the embryos.

### Nuclear-cytoplasmic fractionation and qPCR

Embryos were set on ice and dissociated as described earlier. Nuclear and cytoplasmic fractions were separated by two washes in nuclei isolation buffer (Tris-HCl pH 7.4 10mM, NaCl 10mM, MgCl2 3mM and 0.1% IGEPAL CA-630) followed by 5 minutes centrifugation at 500 x g. RNA was extracted from the two fractions with the RNAeasy Mini Extraction kit (Qiagen, 74044, UK), converted in cDNA via the SuperScript™ III Reverse Transcriptase (Thermo Fisher, 18080093, UK) and used for qPCR.

### Reduced Representation Bisulfite Sequencing (RRBS-Seq)

Genomic DNA were extracted from 4500-7500 sorted somatic cells or PGCs at high, dome and prim-5 stage in duplicates and digested with MspI at 37°C for 3 hours. The fragment ends were repaired with Klenow exo at 37°C for 50 minutes. Then Methylated Illumina Pair-end Adaptors were ligated to gDNA fragments using T4 DNA ligase. The bisulfite conversion were performed using Zymo Research EZ DNA Methylation Gold kit. Libraries were PCR amplified for twenty cycles using Platinum Taq DNA polymerase and sequenced on Illumina HiSeq 2500 on 50bp single-end mode.

### Patch Bisulfite PCR

Genomic DNA were extracted from 5000 sorted somatic cells or PGCs at prim-5, with three replicates for each cell types. Whole Genome Amplified genomic DNA (WGA gDNA) were generated from prim-5 WT Tu fish embryos using GE GenomiPhi V2 DNA Amplification Kits. 30 ng of DNA from sorted cells, 100 ng of genomic DNA from prim-5 embryos (non-bisulfite conversion control) and 100 ng of WGA gDNA (hypomethylation control) were digested with HpyCH4V and NlaIII at 37°C for 1 hour following by heat inactivation for 20 minutes at 65°C. Then custom universal primers were ligated to the targeted fragments using HiFi Taq DNA ligase with the help of gene specific designed oligo patches. The reaction was incubated at 95°C for 15 minutes followed by 30 seconds at 94°C and 4 minutes at 65°C for 25 cycles, and was held at 4°C. Unligated DNA fragments were removed by Exo I and Exo III treatment at 37°C for 1 hour followed by heat inactivation at 80°C for 20 minutes. Bisulfite Conversion were performed following Zymo EZ DNA Methylation Gold kit manufacturer’s instruction. This step was skipped for non-bisulfite control sample. The eluted DNA were PCR amplified using EpiMark Hot Start Taq DNA polymerase. Libraries were sequenced on Miseq 250bp paired-end mode.

### ATAC-seq library preparation and sequencing

Two biological repeats for PGCs and somatic cells at high (wild type), prim-5 (wild type) and morpholino-injected prim-5 (5mm and MO) were prepared. Cells were sorted into 500 µl of cold PBS Mg-, Ca- and immediately treated for ATAC. Nuclear isolation and transposition reaction occurred as described in Buenrostro et al., 2013. Tagmented DNA libraries were purified with 1.2X volume of AMPure XP beads. DNA bound to the beads was washed twice in 80% ethanol and eluted in 20 µl of water. Indexed fragments were checked in concentration by qPCR, profiled by Bioanalyzer, equimolarly pooled and sequenced on an Illumina Next-Seq 550.

### RNA-seq library preparation and sequencing

cDNA was prepared from two biological repeats according to manufacturer’s instructions as follows. Two hundred cells were sorted in 8.5μl of water (0.2U/μl RNase inhibitor). Immediately after collection, cells were added with 1μl of lysis buffer (0.2U/μl RNase inhibitor) and 1 ul of ERCC Mix1-2 (final dilution 1×10^-6^) and flash-frozen. Reverse transcription was performed following the SMART-Seq v4 protocol. Indexed fragments were checked in concentration by qPCR, profiled by Bioanalyzer, equimolarly pooled and sequenced on an Illumina Next-Seq 550.

### ATAC-seq analysis

Paired-end ATAC reads were mapped to the genome using Bowtie2, not allowing discordant mapping of reads (--no-discordant) and insert sizes larger than 5000bp (-- maxins 5000). Results were filtered for mapping quality (10) and for mapping to chromosomes 1 to 25 with the exclusion of the mitochondrial chromosome and of contigs present in danRer7/10 assemblies. There was no removal of subsequent read pairs mapping to the same locus (so called duplicate removal).

Mapped and filtered ATAC reads were corrected for Tn5 transposase overhang by adding 5bp to the position of the start of the first read and by subtracting 4bp from the end of the second read in the read pair as described in Buenrostro et al., 2013. Both thus obtained Tn5 cut sites were extended by a fixed amount. For genome browser visualisation 25bp was added to Tn5 cut sites yielding two 51bp-long regions for each read pair. For PCA and SOM analysis a shorter 5bp extension was used. No selection for particular insert size fraction was used.

### Enhancer calling in somatic cells and PGCs

The set of putative enhancers was obtained from ATAC-seq in the following way. MACS2 was used with options -f BED -g 1.412e9 --keep-dup all --nolambda --nomodel to call peaks in each sample and replicate separately. Four replicate pairs in PGCs and somatic cells at high and at prim5 stages were used to identify peaks reproducible across replicates with Irreproducible Discovery Rate (IDR)2 approach. At 5% IDR the number of reproducible peaks were for soma/high: 12,862; PGC/high: 14,029; soma/prim5: 79,494; PGC/prim5: 61,641. After a further removal of peaks within 500bp from any known transcript start (ENSEMBL version 79/91) and of width greater than 1000bp remaining peak numbers dropped to soma/high: 4,935; PGC/high: 5,097; soma/prim5: 55,784; PGC/prim5: 36,071.

As the last step four peaks sets were merged in a union and thus unified peaks were once again filtered for length <= 1000bp. This yielded a total of 70,612 candidate enhancers.

### Principal Component Analysis and Self Organising Map clustering

In order to assign open chromatin scores to enhancers, windows of size 601bp around enhancer centre were used. ATAC signal levels from genome browser track were extracted (sum of signal values in 601bp bins proportional to the number of 5’ ends of reads falling into these bins and normalised to the total number of reads in each sample) with the help of genomation package3 and saved into a matrix. For PCA and SOM, levels were log-transformed and “centred” by subtracting the mean of each matrix column (sample). For SOM, an additional row (enhancer) centering was performed effectively making SOM operate on log-fold change values.

### Bioinformatic analysis for RRBS

Fastq files were aligned to ZV10 genome and processed using bismark(--bowtie1). Methylation level data were collected using bismark_methylation_extractor with parameters of --bedGraph --cutoff 6 --merge_non_CpG –comprehensive. Following methylation data were analysed using methylKit package in R with three replicates for each cell types at different developmental stages.

### Bioinformatic analysis for Patch bisulfite PCR

Adapters were removed using cutadapt with parameters of -a AGTGTGGGAGGGTAGTTGGTGTT -A ACTCCCCACCTTCCTCATTCTCTAAGACGGTGT -- minimum-length 10 for Read 1 and Read 2. Adapter trimmed fastq files were then aligned to ZV10 genome and processed using bismark (--bowtie2). Methylation level data were collected using bismark_methylation_extractor with parameters of --bedGraph --cutoff 6 --merge_non_CpG –comprehensive. The output CpG coverage files were converted to colorBED files using a custom script. The colorBED files were loaded and visualized on UCSC genome browser.

### RNA-seq analysis

Fastq files were checked for quality by fastqc and trimmed by trimmomatic. Sequencing reads were aligned to the zebrafish genome (danRer9/10) by STAR (v.2.6.1) with the following settings: *--seedSearchStartLmax 12 --outFilterScoreMinOverLread 0.3 -- alignSJoverhangMin 15 --outFilterMismatchNmax 33 --outFilterMatchNminOverLread 0 -- outFilterType BySJout --outSAMunmapped Within --outSAMattributes NH HI AS NM MD -- outSAMstrandField intronMotif --outWigType bedGraph --quantMode GeneCounts*.

Raw read counts were loaded into R and differential expression analysis over samples and stages was performed using DESeq2 (v.1.6.3) and maSigPro packages.

### Statistics

All experiments for which statistical analyses were performed were repeated three times. All sequencing experiments were performed in biological duplicates with the exception of ATAC-seq for dome and 10-somites stages. Data from independent biological repeats were pooled together and the statistical distribution of the dataset was evaluated upon Shapiro-Wilk test. For normally distributed datasets, the p-value was estimated upon t-test, while for non-normally distributed datasets, Wilcoxon test was used.

---

**Supplementary Figure 1.**
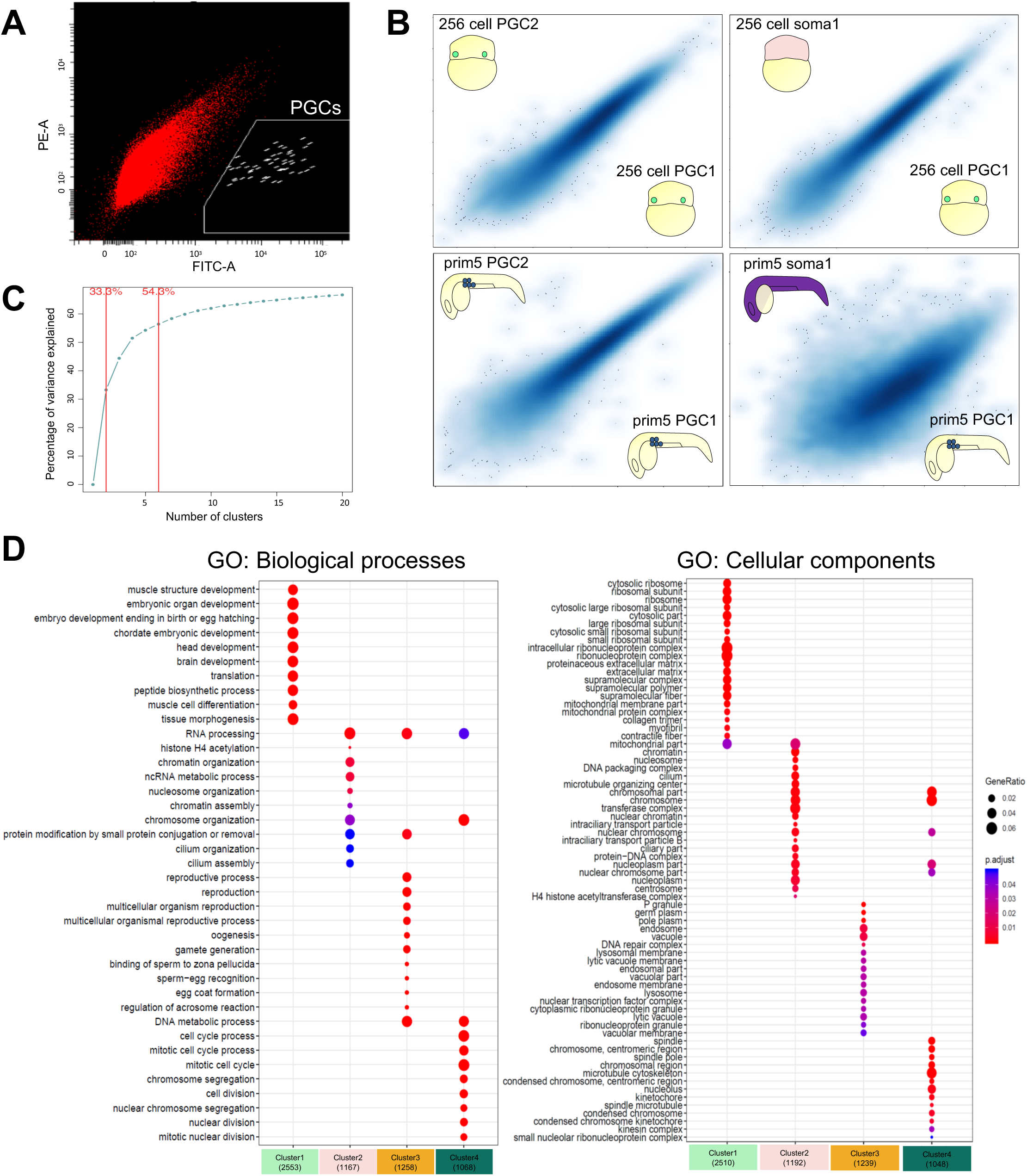
Characterisation of PGC transcriptome highlights early developmental similarities and late divergence between PGCs and somatic cells. (A) FACS profile of GFP-expressing PGCs at prim-5 stage. GFP-positive and –negative cells are in white and red respectively. (B) Transcriptome correlation profiles after RNA-seq reads normalisation. Read counts for each gene were normalised for library size and gene length (tpm) and correlation among stages and cell types were compared. Reads are shown as Log_2_(tpm+1). (C) Justification of number of k-means selected based on inter-sample variance. (D) Biological processes and cellular components GO analysis for the four identified expression groups by k-mean clustering (p adjusted < 0.05).

**Supplementary Figure 2.**
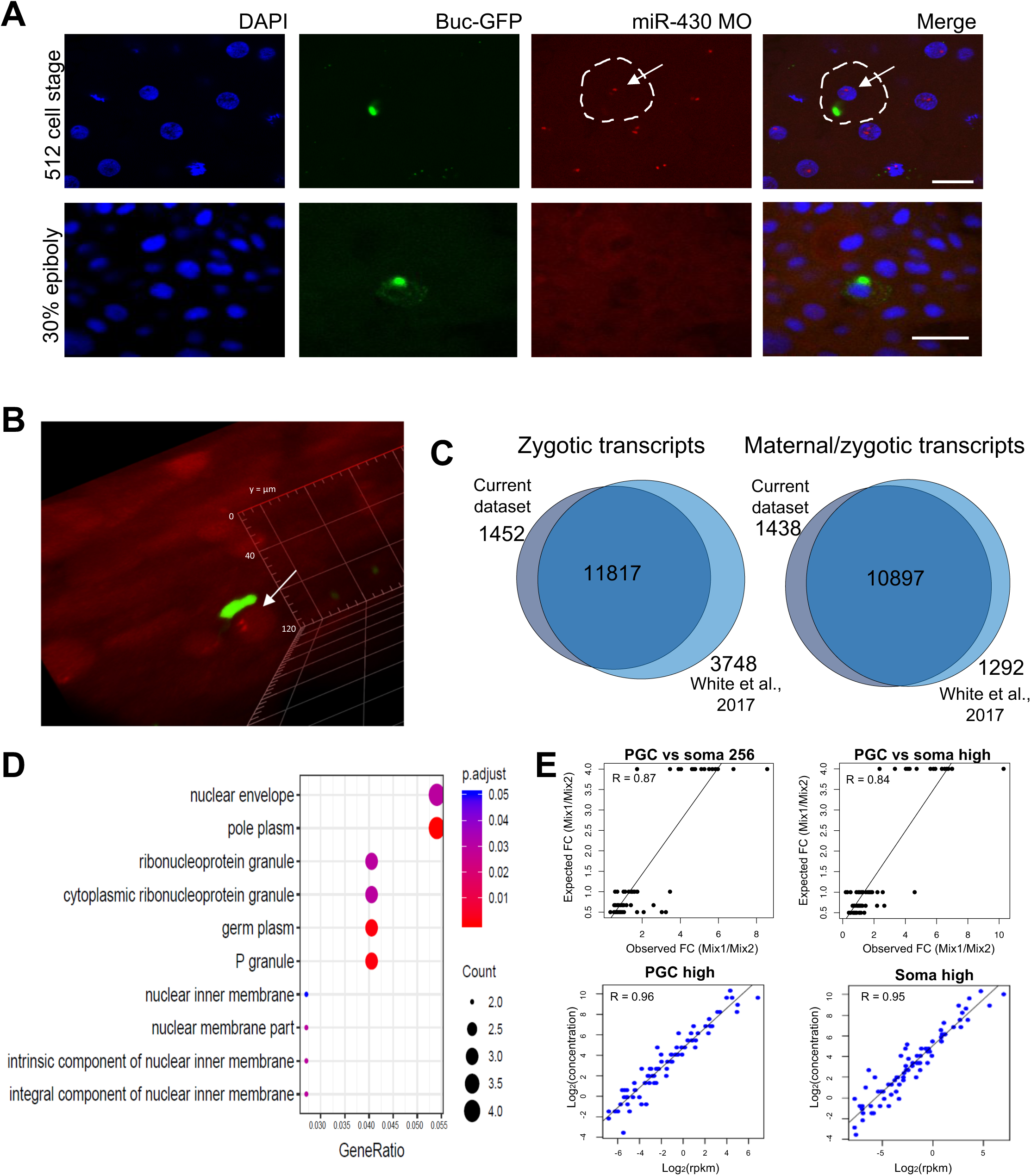
PGCs do not delay transcriptional activation. (A) Maximum intensity projection of a multi-stack image of *miR-430* transcriptional activity at 512-cell stage and 30% epiboly. PGC boundaries are highlighted as white, dashed lines. Arrow indicates the nucleus of a germ plasm-carrying cell where transcriptional foci are detected. Scale bar is 30 µm. (B) 3D rendering of a *miR-430-*expressing nucleus (arrow) and a GFP-tagged germ granule. White arrow indicates the germ cell nucleus. (C) Venn diagram for predicted zygotic/maternal and zygotic genes from two independent RNA-seq datasets. (D) Cellular components GO analysis for genes upregulated in PGCs after the first wave of ZGA (from 256-cell to high stage). P adjusted < 0.05. (E) Representative regression scatter plots between expected vs observed ERCC mix ratios (top) and ERCC concentrations vs ERCC reads (bottom). The tpm threshold for normalised ERCC reads was 1.

**Supplementary Figure 3.**
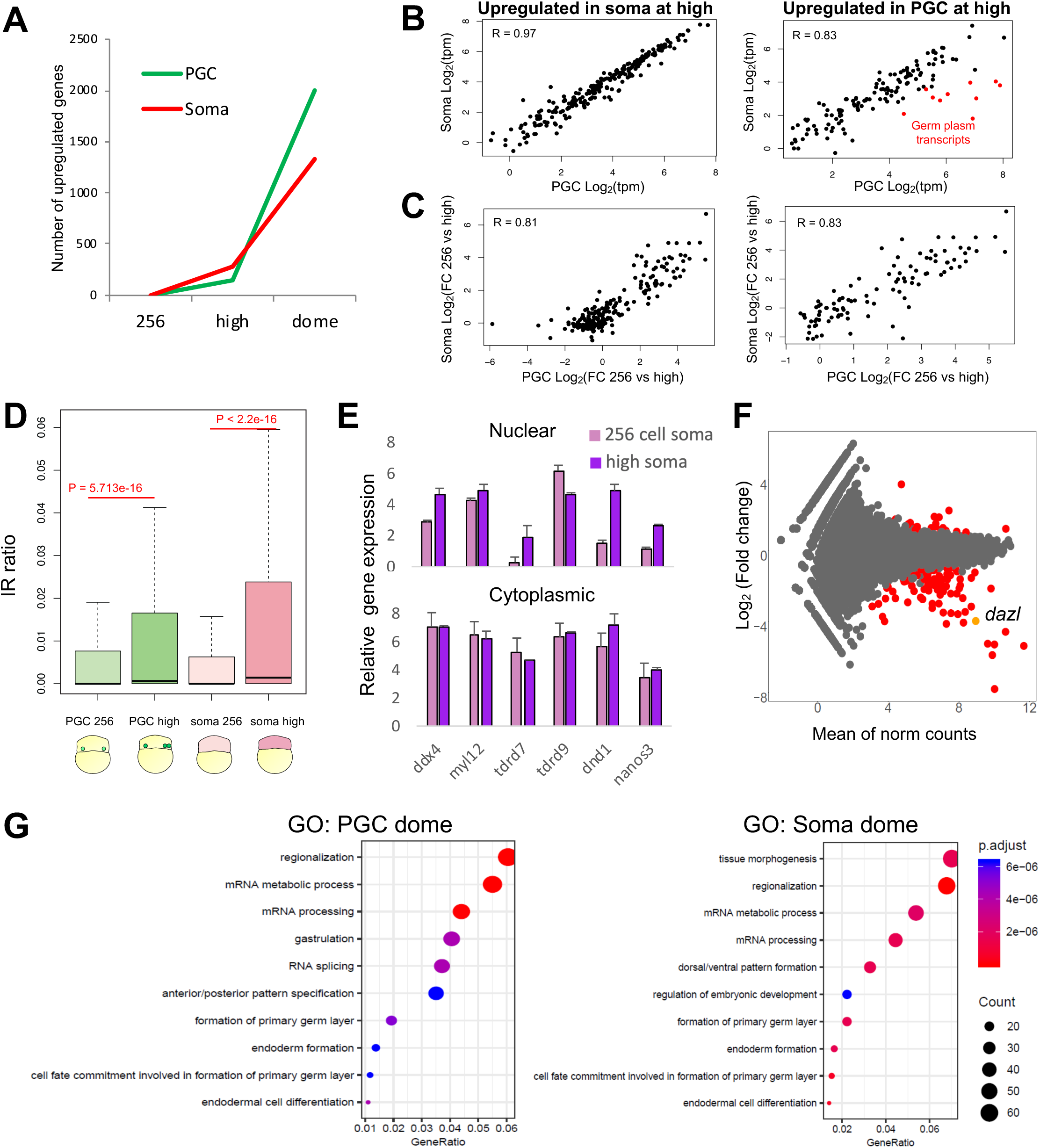
Differential transcriptome between PGCs and somatic cells at early stages is not caused by differential transcription. (A) Count of upregulated genes from previous stage in PGCs and somatic cells at high and dome stages. (B) Correlation of normalised RNA-seq reads (tpm) for genes upregulated from the previous stage in the PGCs or somatic cells only at high stage. Germ plasm genes are in red. (C) Correlation between fold change increases from 256-cell to high stages in PGCs and somatic cells for gene subsets. (D) Intron retention ratio for all transcripts in PGCs (green) and somatic cells (purple) before and after ZGA. T-test was used to calculate p-values. (E) Relative gene expression fold change normalised to a reference gene in somatic cells for nuclear and cytoplasmic fractions. Error bars show standard errors. (F) Differential expressed genes between 256-cell and high stages in PGCs. Significantly differentially expressed genes are in red. P adjusted < 0.1, log_2_FC < −1/> 1. (G) Biological processes GO analysis for genes upregulated from high to dome in PGCs (left) and somatic cells (right).

**Supplementary Figure 4.**
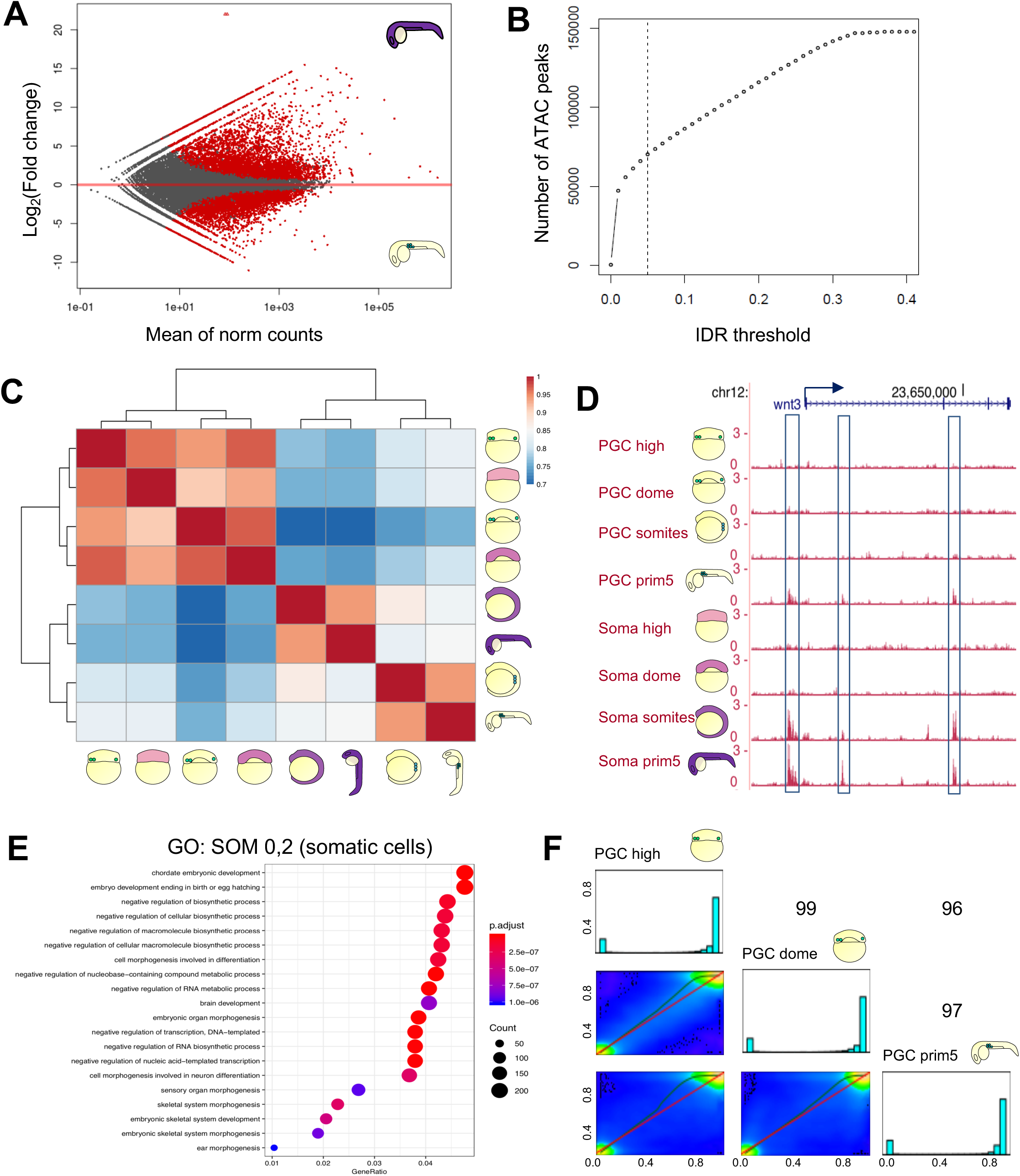
Gradual acquisition of germ identity is accompanied by epigenetic changes. (A) Differentially expressed genes in PGCs and somatic cells at prim-5 stage. Significantly differentially expressed genes are in red. P adjusted < 0.1. (B) Reproducible ATAC peaks identified in function of the IDR threshold. (C) Unsupervised correlation heatmap for developmental stages and cell types after selection of reproducible ATAC peaks. (D) Genome browser view showing ATAC-seq tracks (magenta) of *wnt3* gene in PGCs and somatic cells across early development. Blue boxes highlight putative regulatory elements. Arrows show transcriptional directionality. (E) GO terms for genes associated with open chromatin regions upregulated in late somatic cells. (F) CpG base Pearson correlation shown as heatmap and histogram of CpG frequency.

**Supplementary Figure 5.**
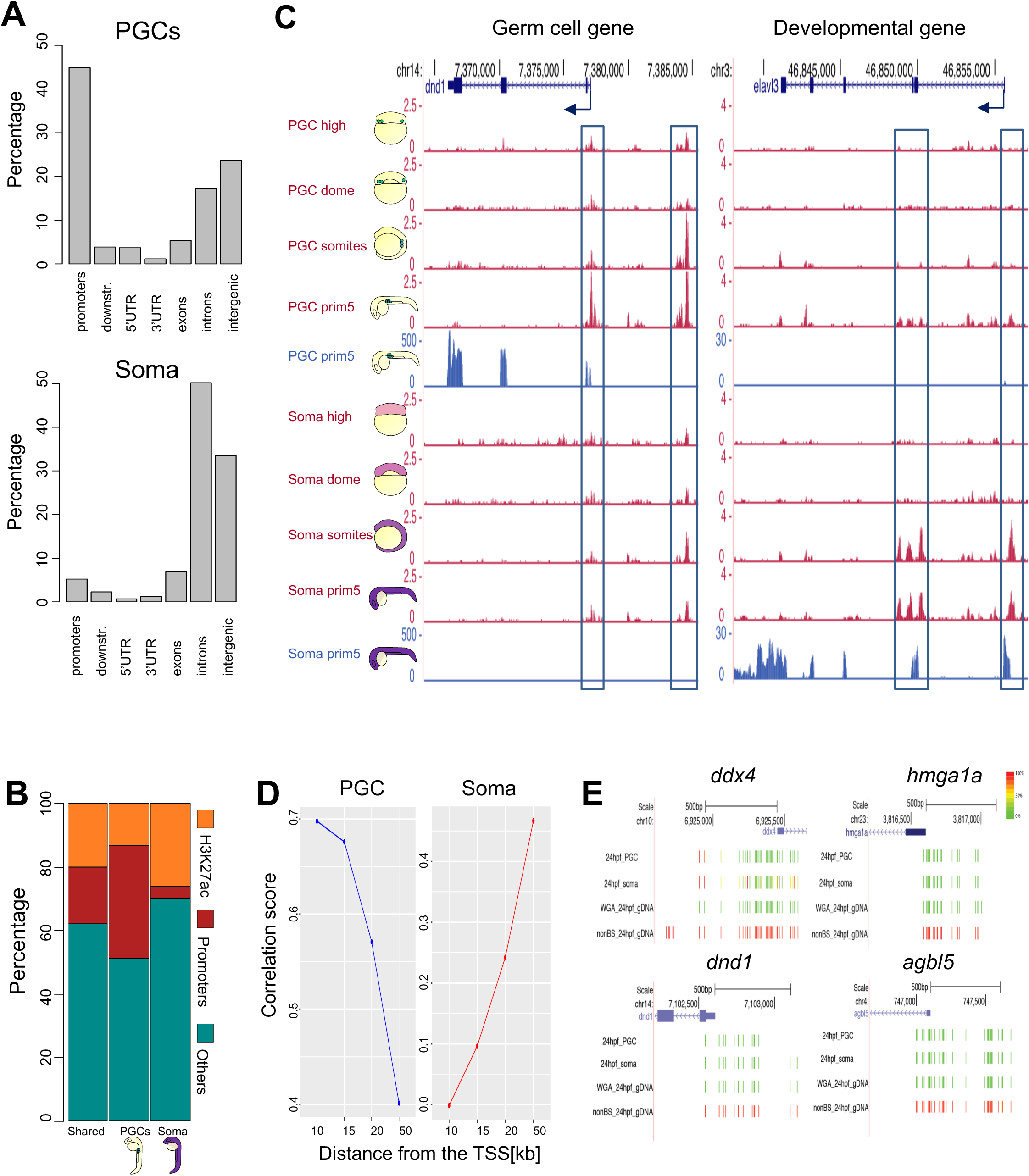
PGCs do not open chromatin at regions identified as putative enhancers. (A) Percentage of differentially accessible ATAC peaks in PGCs and somatic cells at prim-5 stage overlapping gene features. Promoters include 1kb up- and downstream from the TSS. (B) Genome browser view of open chromatin regions (ATAC-seq in magenta) and transcript levels (RNA-seq in blue) in PGCs and somatic cells. Blue boxes highlight putative regulatory elements or promoters. Arrows show transcriptional directionality. (C) Percentage of ATAC peaks associated with H3K27ac and promoters in PGCs and somatic cells at prim-5 stage. (D) Correlation score between chromatin accessibility and gene transcription in function of distance from the TSS. Correlation score is represented as the absolute ATAC log2FC value associated to genes upregulated in PGCs (blue) or somatic cells (red) (E) Genome browser view of four representative promoters upregulated in late PGCs versus somatic cells. CpG methylation levels are reported as coloured bars.

**Supplementary Figure 6.**
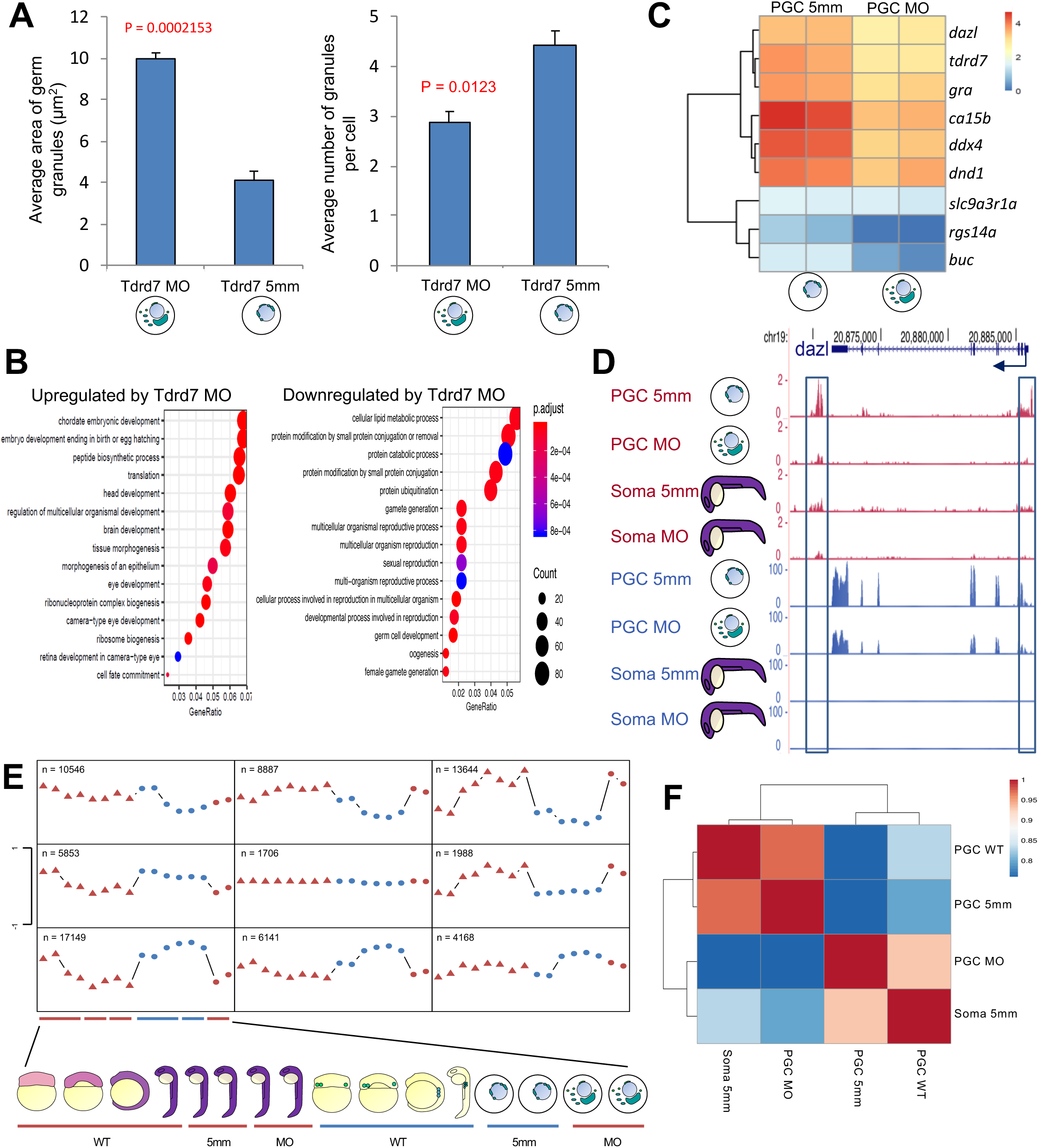
Tdrd7 translation is required for maintaining PGC specification. (A) Phenotypic effect of Tdrd7 KD PGCs. Number and size of germ granules were measured. P-value against control is shown. Red colour indicates significant difference (p < 0.005) based on Wilcoxon test. (B) GO terms for genes upregulated and downregulated by Tdrd7 translational inhibition in PGCs. (C) Gene expression heatmap for germ plasm-localised transcripts. Colour intensities indicate log_2_(tpm+1). (D) Genome browser view of ATAC profiles after morpholino injections. Open chromatin (ATAC-seq) is shown in magenta and normalised transcript levels (RNA-seq) are shown in blue. Arrows show transcriptional directionality. (E) Self-organising map of reproducible open chromatin regions. Wild type (WT) PGCs and somatic cells are represented as circles and triangles respectively. Tdrd7 KD PGCs are shown as red circles. (F) Unsupervised correlation heatmap between wild type (WT) mismatch- and *tdrd7-*targeting morpholinos.

